# Structural mechanism of nuclear membrane sealing by LEM2-ESCRT-III

**DOI:** 10.64898/2026.09.01.748678

**Authors:** Mareike A. Jordan, Karen Palacio-Rodriguez, Ramesh Adakkattil, Sonja Welsch, Lukas Niese, Dollie LaJoie, Esther von der Weth, Arthur Alves Melo, Lisa Redlingshöfer, Shotaro Otsuka, Stefan Diez, Gerhard Hummer, Alexander von Appen

**Affiliations:** Max Planck Institute of Molecular Cell Biology and Genetics; Pfotenhauerstraße 108, 01307 Dresden, Germany; Department of Theoretical Biophysics, Max Planck Institute of Biophysics, 60438 Frankfurt, Germany; Central Electron Microscopy Facility, Max Planck Institute of Biophysics, Max-von-Laue-Straße 3, 60438 Frankfurt am Main, Germany; B CUBE – Center for Molecular Bioengineering, TUD Dresden University of Technology, 01307 Dresden, Germany; Human Technopole (HT), Molecular Cell Biology, Viale Rita Levi-Montalcini, 1 - Area MIND - 20157 Milano Italy; Institute for Neurodegenerative Diseases, University of California San Francisco, San Francisco, California 94158, United States; Max Perutz Labs, Vienna Biocenter Campus (VBC), Vienna, Austria; Medical University of Vienna, Max Perutz Labs, Vienna, Austria; Department of Theoretical Biophysics, Max Planck Institute of Biophysics, Goethe University Frankfurt, 60438 Frankfurt, Germany

## Abstract

In open mitosis, re-establishing nucleocytoplasmic compartmentalization requires the LEM2-ESCRT machinery to coordinate spindle clearance with sealing of the remaining nuclear envelope pores. The structural basis of this topologically unique and fundamental membrane-remodeling process is poorly understood. Here, we combine biochemical reconstitution, cryo-electron tomography, subtomogram averaging and large-scale molecular dynamics simulations to define the structural mechanism of nuclear membrane sealing. We structurally resolve that LEM2’s winged-helix domain (WH) co-polymerizes with the ESCRT-II/III protein CHMP7 to form a membrane-bound scaffold whose geometry is progressively remodeled by downstream ESCRT-III proteins as it transitions from the flat membrane surrounding the pore towards the negatively curved membrane neck. In parallel, LEM2 positions its intrinsically disordered low-complexity domain within the pore, where condensation around spindle microtubules mechanically couples the membrane-ESCRT-LEM2 scaffold to the spindle and narrows the remaining diffusion path, restoring compartmentalization before membrane closure is complete. Remarkably, the LEM2-WH domain alone forms tightly constricted membrane tubes, coating the negatively curved inner surface, revealing an intrinsic membrane-remodeling activity of the receptor itself. Together, our work establishes a structural framework for how receptor-ESCRT co-polymerization, low complexity domain-mediated sealing and receptor-driven membrane remodeling guide nuclear-envelope pores from spindle-containing openings to terminal constriction and fusion.

## Main Text

Cellular function depends on compartmentalization of the genome by the nuclear envelope (NE), a double-membrane system that separates chromatin from the cytoplasm. During open mitosis, this compartment is transiently dismantled to allow chromosome segregation and must be rapidly re-established during mitotic exit. The NE reforms from endoplasmic reticulum-derived membranes that spread over the chromatin surface (1,2). However, spindle microtubules remain attached to chromatin until late anaphase, creating an obstacle within catenoid-shaped double-membrane pores (hereafter, membrane pore) that must be sealed to restore nuclear function (3) (Fig. 1a). The nucleus regains nucleocytoplasmic compartmentalization even before spindle disassembly, which creates a unique topological challenge: to form a diffusion barrier to the cytosol while rigid microtubules still cross the remaining membrane pore.

**Figure 1:**
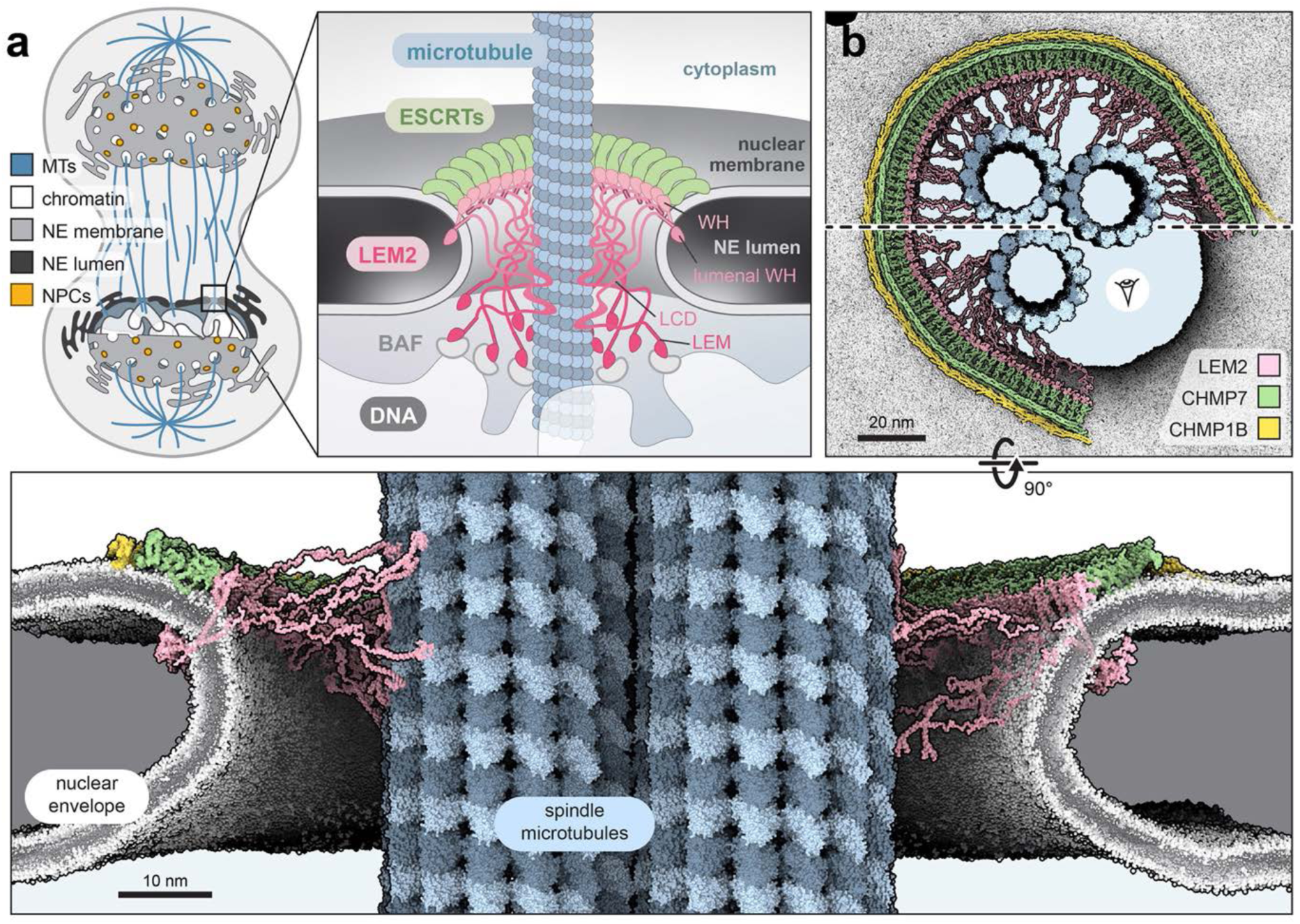
LEM2 coordinates membrane remodeling and microtubule sealing during nuclear-envelope closure. **a)** Cartoon model: At the end of mitosis, the NE closes around spindle MTs that are still attached to chromatin, forming catenoid membrane pores, that must be sealed to re-establish nucleocytoplasmic compartmentalization and preserve nuclear integrity. Within these pores, LEM2 creates the interface between membrane, ESCRT-III and microtubules at the chromatin surface. **b)** Structural model of LEM2-ESCRT-III-mediated sealing of a spindle-containing NE pore. The model depicts a short-lived intermediate after 200 ns of MD simulation and is based on the experimentally determined structure of the LEM2_WH_-CHMP7-CHMP1B scaffold and the geometry of spindle-containing membrane pores observed in cells.

Genetic and cell biological studies identified the conserved LEM2-ESCRT pathway as the machinery responsible for this process (4–8). The inner nuclear membrane protein LEM2 recruits the ESCRT-II/III hybrid CHMP7 to spindle microtubules (6,8), which subsequently assembles downstream ESCRT-III proteins, including CHMP4, CHMP2, CHMP1B and Ist1 at closing membrane pores (4,5). ESCRT-III proteins form dynamic filaments that transition between distinct architectures in response to filament composition and regulation (9–15). At the NE, ESCRT activity is orchestrated by the transmembrane protein LEM2, which coordinates the key molecular interfaces required for NE sealing: while inserted into the inner nuclear membrane, its winged-helix domain binds CHMP7, its LEM domain binds the chromatin adaptor Barrier-to-autointegration factor (BAF), and its intrinsically disordered low-complexity domain (LCD) condenses upon the surface of spindle microtubules and undergoes condensation (4–6,8) (Fig. 1a). These observations led to a model in which LEM2 and ESCRT-III form a molecular O-ring around spindle microtubules, transiently restoring compartmentalization while membrane closure remains incomplete (8). This principle has been shown to be conserved in yeast (16). Yet how ESCRT filaments shape nuclear membrane pore geometry while LEM2 seals the residual space around spindle microtubules and how these processes couple to terminal membrane constriction remains unresolved.

Here, we determine the structural mechanism of LEM2-ESCRTIII-mediated nuclear membrane sealing. We show how LEM2 successively regulates ESCRT polymerization and membrane remodeling while deploying its disordered domains within the pore to restore nuclear compartmentalization after mitosis (Fig. 1b).

## Results

### The nuclear sealing reaction can adapt to diverse pore geometries

Nuclear membrane sealing occurs on the minute timescale during mitotic exit going from large to small openings around spindle microtubules. To define the membrane topology encountered by the LEM2-ESCRT machinery when the machinery is active (4,5,8), we analyzed segmentations of room-temperature tomograms from high-pressure frozen anaphase HeLa cells (1,17). We observed catenoid-shaped membrane pores in the double membrane of the NE resembling the membrane shapes of nuclear pore complex (NPC) assembly intermediates, but enclosing spindle MTs that traversed perpendicularly through the membrane openings into the nucleus, consistent with previous observations (3) (Fig. 2a, Supp. Fig.1a,b). Compared with NPC assembly intermediates, spindle microtubule enclosure sites varied substantially in size, containing between 1 and more than 20 microtubules (Supp. Fig. 1c, d). Thus, the nuclear membrane sealing machinery must adapt to a broad range of pore geometries. Despite this variability, most enclosure events contained only one single MT. Rotational averaging revealed a continuous ring-like density occupying the narrow space between the microtubule surface and the surrounding membrane (Fig. 2a, Supp. 1Fig. c,d), at a stage when LEM2-ESCRT components are known to accumulate around spindle MTs (4–6,8). Notably, we did not observe membrane openings devoid of either spindle microtubules or NPCs consistent with the model that membrane pores close by themselves (18). Only a single tomogram captured microtubules immediately after detachment from chromatin, leaving a small membrane opening of ∼50 nm and a condensed chromatin surface underneath (Supp. Fig. 2), indicating that spindle release and pore closure occur within a narrow temporal window that is difficult to capture within in cells. Together, these observations define the geometric constraints of nuclear membrane sealing: despite variation in pore size and microtubule number, the LEM2-ESCRT machinery encounters a common topology, a membrane pore surrounding one or more spindle microtubules. Because these intermediates are inherently short-lived in cells, we turned to biochemical reconstitution to stabilize and structurally dissect the underlying LEM2-ESCRT assemblies.

**Figure 2:**
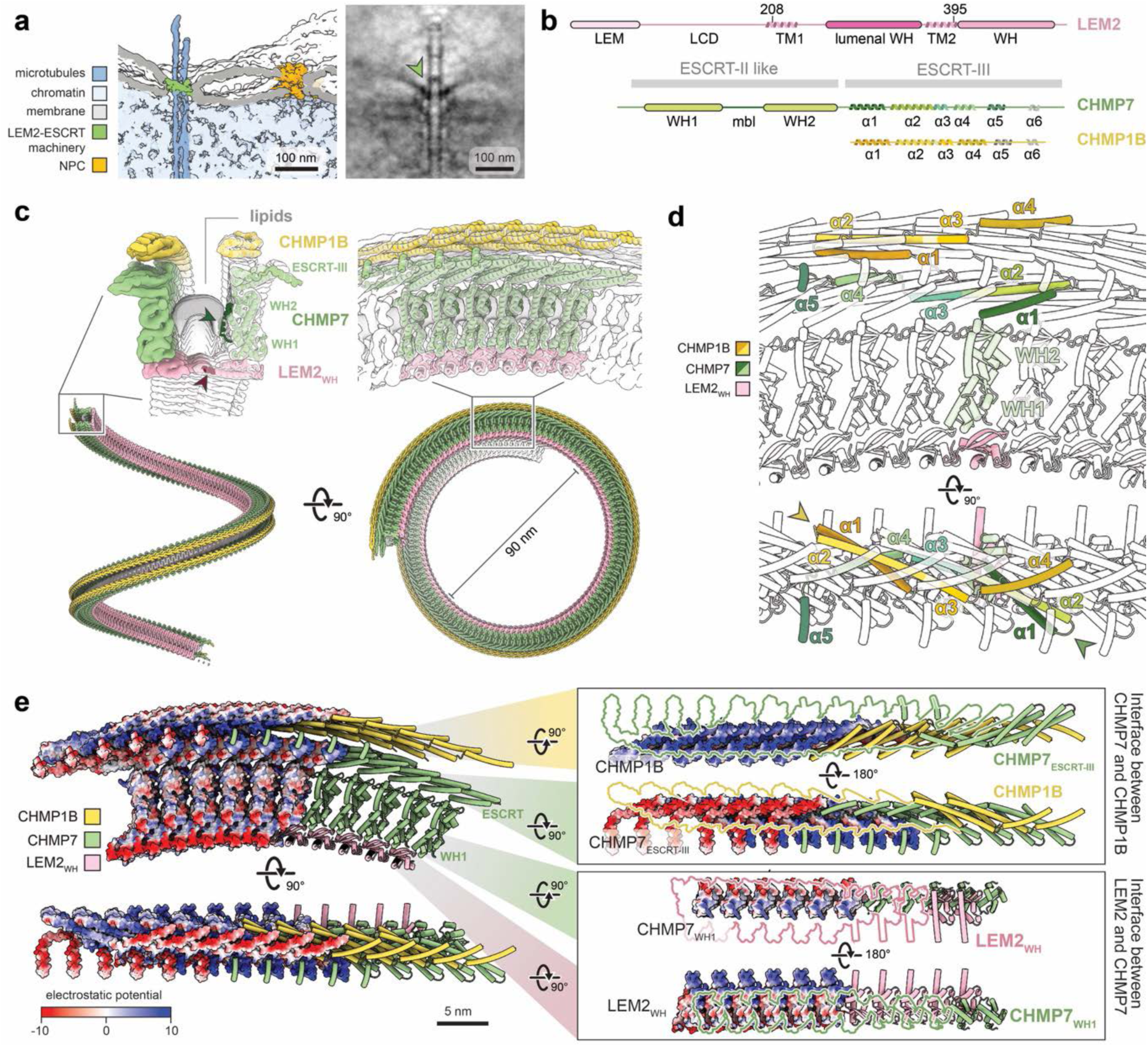
High-resolution structure reveals that LEM2 is an integral component of the ESCRT-III polymer. **a)** In-cell topology of a closing membrane pore around a spindle MT, 7.7 min after anaphase onset in a HeLa cell; left: segmentation; right: slice of a rotational average. **b)** Domain organization maps of proteins used in this study. **c)** cryo-ET subtomogram average and modeled structures of LEM2_WH_, CHMP7 and CHMP1B from C2-symmetric filament reconstitutions with solubilized lipids. Arrow heads: CHMP7 membrane-binding-loop (mbl) and LEM2_WH_ lipid-anchoring residues. **d)** Subunit placement within the filament. ESCRT-III helices are labeled. **e)** Electrostatic surface of the LEM2_WH_-CHMP7-CHMP1B filament. Right: flipped-open view of strand interfaces. The respective interaction partners have been removed from the views and are indicated as colored outlines. The reconstruction was obtained from 38,278 subtomograms at an estimated global resolution of 7.3 Å using the FSC = 0.143 criterion (Supp. Fig. 2, Methods).

### LEM2 co-polymerizes with ESCRT-III

To determine how the LEM2-ESCRT machinery assembles during membrane sealing, we reconstituted the ESCRT receptor winged helix (WH) of LEM2 with full-length CHMP7, with or without the downstream ESCRT-III protein CHMP1B (Fig. 2b), using solubilized lipids and dialysis (19). Consistent with previous observations (8), LEM2_WH_ induced the processive polymerization of CHMP7 into large-curvature filaments. Cryo-ET revealed characteristic curvatures of 90-120 nm diameter, ranging from ∼40 to 200 nm (Supp. Fig. 3), matching the size range of membrane openings observed around microtubule bundles in anaphase cells (Supp. Fig. 1a,b). Subtomogram averaging yielded a 7.3 Å reconstruction of the LEM2_WH_-CHMP7-CHMP1B filament (Fig. 2c, Supp. Fig. 4) with a two-stranded antiparallel C2 architecture, indicating that filaments can polymerize in both directions. Interestingly, LEM2_WH_, CHMP7 and CHMP1B appeared iso-stoichiometrically, revealing a co-polymerization mechanism. LEM2_WH_ occupies the inner arc of the curved filament, where each subunit interacts with two WH1 domains of CHMP7, thereby driving cooperative filament assembly through alternating lateral stabilization (Fig. 2c,d). The progeria-associated mutation LEM2S479F (20) maps directly to this interface (Supp. Fig. 5a). The addition of the late ESCRT-III protein CHMP1B to LEM2_WH_-CHMP7 stabilized the filaments into more regular assemblies (Supp. Fig. 3) better suited for subtomogram averaging. In the absence of LEM2_WH_, CHMP7 failed to polymerize processively (Supp Fig. 6). However, consistent with previous results (8) CHMP7 assembles into flexible spirals in the presence of membranes. This suggests that LEM2 regulates CHMP7 polymer diameter and stability. Taken together, our structure reveals that LEM2 is not merely an ESCRT recruiter but an integral, stoichiometric component of the ESCRT scaffold itself. LEM2_WH_ recruits the ESCRT-III machinery to sites of nuclear fusion via the ESCRT-II-like CHMP7_WH1_ adaptor domain, positioning CHMP7_ESCRT-III_ for cooperative polymerization into defined geometries.

### ESCRT-III opening enables co-polymerization and tunes filament mechanics

Previous work suggests that conserved ESCRT-III helices adopt closed or open conformations, with the open state characterized by extension of α2 into a continuous α2-α3 helix (21–23). In the LEM2_WH_-CHMP7-CHMP1B filament, both CHMP7 and CHMP1B adopt an open conformation (Fig. 2d). Previous XL-MS data (8) and AlphaFold predictions indicate that CHMP7 is closed in solution. Thus, filament assembly proceeds by opening of the ESCRT-III, enabling each subunit to establish an interwoven network with four additional subunits (i+4 polymerization) as previously described for CHMP1B (21) (Supp. Fig. 7a). Consistent with this mechanism, all-atom MD simulations showed that the open conformations of CHMP7 and CHMP1B are stable only within the membrane-bound polymer, whereas isolated monomers of these proteins revert toward the closed state in solution (Supp. Fig. 7b-d). Thus, filament assembly stabilizes the open ESCRT-III conformation and generates an interwoven network of lateral contacts that enables cooperative co-polymerization. As observed for other ESCRT-III copolymers (21,24,25), CHMP7 and CHMP1B interact predominantly through electrostatic complementary: an elongated acidic surface formed by CHMP1B-helix 1 engages a corresponding basic surface formed by CHMP7_ESCRT-III_-helix 4 (Fig. 2e). Given that LEM2_WH_-CHMP7 filaments assembled in the absence of CHMP1B retained the same overall architecture but displayed a broader curvature distribution (Supp. Fig. 3), our data suggests that CHMP1B hence stabilizes filament geometry changing preferred curvature and hence mechanics. Together, these observations suggest a model where LEM2-ESCRT-III copolymerization results in geometrically defined polymers. LEM2-activated CHMP7 forms a charged interface through its interwoven ESCRT-III folds as platform for downstream ESCRT-III recruitment, while CHMP1B association further mechanically constrains the filament curvature, suggesting a rigidification. We therefore asked how these composition-dependent properties influence the response of the filament to membrane-pore geometry.

### LEM2-ESCRT-III forms a single-stranded membrane-bound polymer

The cryo-ET structure of reconstituted LEM2_WH_-CHMP7-CHMP1B filaments revealed an electron density which can be attributed to bicellar lipids (Supp. Fig. 5b), suggesting a defined membrane interface formed by the double-stranded co-polymer. However, the cellular cytoplasm-chromatin asymmetry suggests that the C2 symmetry of the double-stranded co-polymer depicts a reconstitution artefact arising from the truncation of LEM2 transmembrane domains and the use of detergent solubilized lipids (Fig. 2c, Supp. Fig. 5b,c). To confirm the membrane binding interface, we reconstituted LEM2_WH_, CHMP7 and CHMP1B on continuous membranes and analyzed the sample with segmentation and subtomogram averaging. We found, that in contrast to solubilized lipids, preformed membranes of vesicles yielded single-stranded LEM2_WH_-CHMP7-CHMP1B co-polymers that followed membrane curvature as flat spirals or wrapped around vesicles, confirming the membrane binding interface (Fig. 3a,b). Hence, we extended the truncated LEM2_WH_ domain by the transmembrane and lumenal domains as obtained from AlphaFold predictions (Supp. Fig. 5d). We placed a trimeric repeat of this co-polymer in a lipid membrane previously equilibrated at coarse-grained Martini resolution (see Methods) to perform all-atom MD simulations. In these simulations, the structure engages lipid membranes readily and remains stable throughout the entire duration of 1 µs (Fig. 3c). All three scaffold components have interactions with the membrane with an interface that is consistent with the orientation seen in the experiments (Fig. 3b). In addition to its transmembrane helices, a basic cluster in the WH domain of LEM2 (Arg455, Arg456, Arg459, Arg463) frequently contacted phospholipid headgroups. CHMP7 engaged the membrane through its previously described membrane-binding loop between WH1 and WH2 (26), with Leu135 inserting deeply into the hydrophobic core of the bilayer, while Lys337 at the C-terminal end of ESCRT-III helix 3 formed additional lipid contacts. By contrast, the CHMP1B membrane-binding interface previously identified in positively curved CHMP1B-IST1 tubes (27) is buried within the CHMP1B-CHMP7 interface of our structure (Supp. Fig. 8, see discussion). Instead, CHMP1B contacts the membrane through residues between Lys30 and Lys59, adjacent to the loop connecting helices 1 and 2 (Fig. 3c, Supp. Fig. 8). Together, these observations suggest that all components of a single-stranded copolymer directly bind membranes.

**Figure 3:**
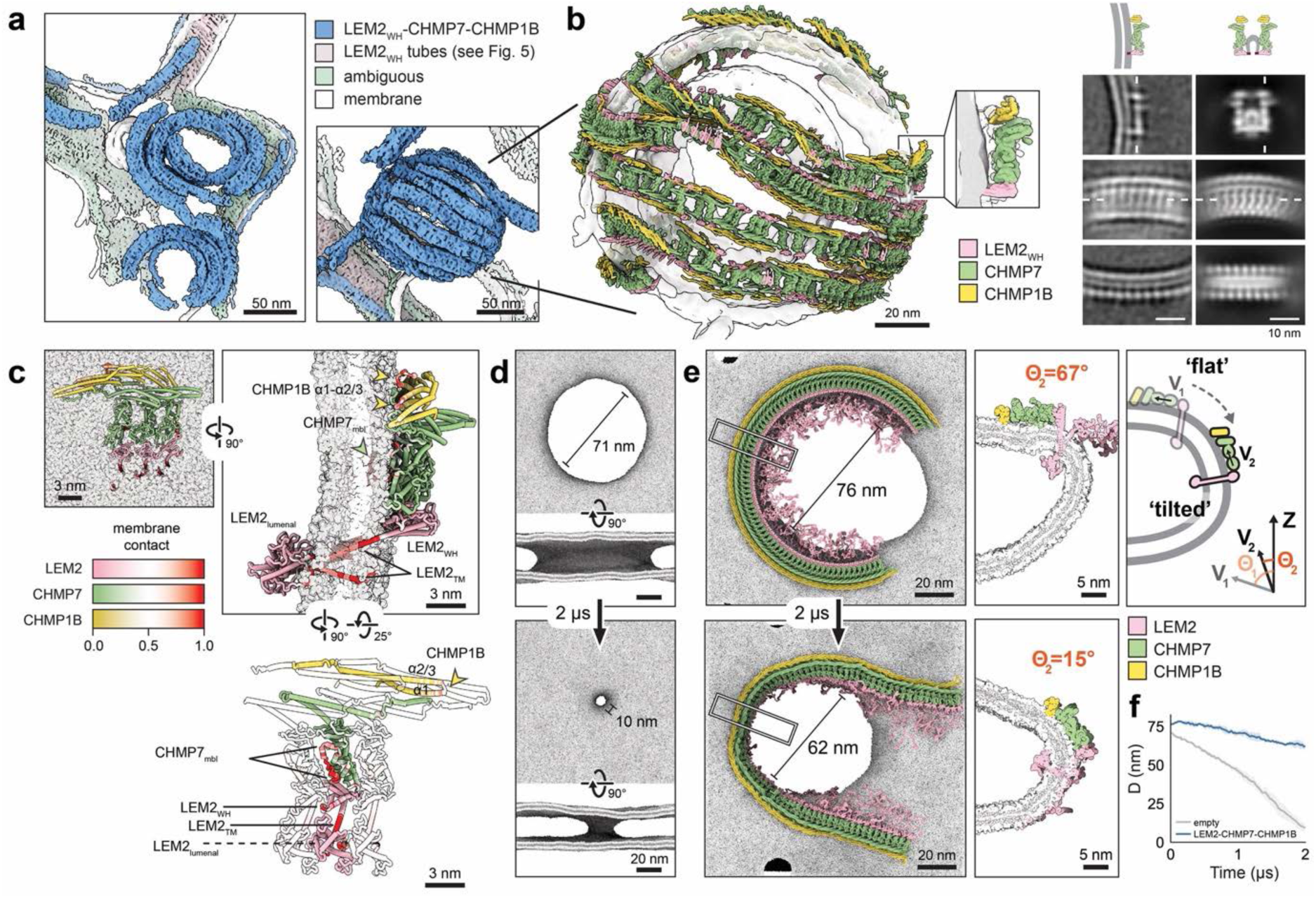
A membrane-bound LEM2-ESCRT-III scaffold stabilizes membrane pores by adapting to their geometry. **a)** Tomographic segmentations of LEM2_WH_, CHMP7 and CHMP1B assembled on vesicles as single-stranded filaments. b) The structure of LEM2_WH_-CHMP7-CHMP1B determined in solubilized lipids was mapped back onto particle positions identified on continuous membranes. Right: Class averages from membrane-bound reconstitutions (left) and solubilized lipids (right) reveal the single-stranded and double-stranded architectures, respectively. c) All-atom MD simulations of a LEM2_ΔLCD_-CHMP7-CHMP1B trimer on flat membrane after 0.5 µs. LEM2 was extended with its transmembrane domains as obtained from AlphaFold predictions. Protein-membrane interactions are mapped in red. d) Coarse-grained MD simulation of a 71-nm membrane pore. e) Coarse-grained MD simulations of a single-stranded LEM2-CHMP7-CHMP1B filament placed on a membrane pore. The filament stabilizes the diameter while its subunits progressively reorient relative to the membrane normal, adapting to the catenoid geometry of the pore neck. The tilt Θ was measured as the angle between the z-axis and vector V connecting the CHMP7 WH1 and WH2 domains. f) Diameter of an empty membrane pore and of a pore containing LEM2-CHMP7-CHMP1B filament as functions of time. N=3 independent simulations were performed per condition; traces show mean values ± standard deviation.

### Conformational changes of LEM2-ESCRT-III in membrane pores

During NE closure, the LEM2-ESCRT-III scaffold must adapt from the flat membrane surrounding the pore to the negatively curved membrane neck of the pore channel. To determine the dynamic interplay of membrane pores and the LEM2-ESCRT-III filament, we performed coarse-grained MD simulations. As a reference, we first simulated an empty ∼70-nm membrane pore with NPC-like geometry. In the absence of protein, the pore rapidly constricted to ∼10 nm within 2 µs (Fig. 3d,f), consistent with previous observations (18). We then placed a LEM2-CHMP7-CHMP1B filament onto the same pore geometry. Given that ESCRT components are actively removed by Vps4 (28,29), we modeled an open filament (a three-quarter-turn ring) to allow flexible adaption (Supp. Fig. 9) and performed MD simulations for 2 µs. We found, that in the presence of the filament, pore closure slowed to ∼62 nm after 2 µs (Fig. 3e,f). Thus, instead of simply promoting pore constriction, the LEM2-ESCRT-III scaffold stabilizes an intermediate open-pore geometry. Rather than remaining static, the scaffold underwent a pronounced conformational transition. The filament center migrated toward the pore channel, converting its initially flat into a tilted conformation. Specifically, the flat-to-tilted conformational change of the filament happened through a rotation of the protein subunits relative to the helical axis, producing a partial inside-out movement of the filament (Fig. 3e, Supp. Fig. 10a). Hence, our simulations are consistent with the theoretically proposed buckling mechanism of ESCRT-III filaments in context of membrane remodeling (30,31). The tilting of the filament center came at the cost of the filament termini moving out of the membrane pore. However, when expanding the membrane pore size with external tension in the MD simulations, the entire filament strand migrated to the tilted conformation, confirming its preferred position at the membrane neck (Supp. Fig. 11a). In conclusion, LEM2-CHMP7-CHMP1B dynamically stabilizes membrane pores and adapts a tilted conformation at the pore membrane neck.

The confirmational changes of the filament must be reflected on the protein component level and hence we asked how the individual components contribute to membrane pore stabilization. Our previous cryo-ET analysis showed that LEM2-CHMP7 form stable but more flexible filaments (Supp. Fig. 3). Concurrently in membrane pore simulations, removing CHMP1B resulted in stable LEM2-CHMP7 filaments, which however, completely remained in their flat conformation with a similar effect on the pore closure dynamics (Supp. Fig. 10b). Instead of tilting, these filaments responded to pore constriction predominantly by increased in-plane bending. Thus, incorporation of a second ESCRT-III strand through CHMP1B increases filament rigidity and constrains in-plane bending. Consequently, membrane pore closure drives the bending of the filament out of its original flat plane, resulting in filament tilting and migration toward the membrane neck. Conversely, simulations of a hypothetical LEM2-only filament within a ∼70-nm membrane pore did not retain its large curvature (Supp. Fig. 11b). This demonstrates that CHMP7 is required to stabilize the large curvature of the early LEM2-CHMP7 filament, consistent with the flat spirals observed experimentally for CHMP7 on membranes (Supp. Fig. 6).

Together, these analyses identify the LEM2-ESCRT-III scaffold as a compositionally tunable polymer whose mechanical properties determine how it adapts to membrane-pore geometry. CHMP1B constrains in-plane bending, redirecting deformation into filament buckling and thereby positioning the scaffold within the membrane neck. This positions the scaffold at the site where the forming NE membrane encounters spindle microtubules, which themselves are obstacles to membrane closure. We therefore asked how the LEM2-ESCRT-III scaffold couples to microtubules within the pore.

### LEM2_LCD_ couples the membrane scaffold to spindle microtubules

To determine how the LEM2-ESCRT-III scaffold engages spindle microtubules during NE sealing, we reconstituted membrane-bound LEM2_LCD_-His_6_ on Ni-NTA-containing (4 %) vesicles, added stabilized microtubules and analyzed by cryo-ET. Consistent with previous work (8), LEM2_LCD_-His_6_ condensed around microtubules, bundled them and tethered membranes tightly to the microtubule surface (Fig. 4a, Supp. Fig. 12a-c). LEM2_LCD_-His_6_ additionally induced vesicle blebbing and cargo-driven intraluminal vesicle formation (Supp. Fig. 12d,e), revealing that condensation of the LEM2_LCD_ is sufficient to deform membranes, in line with recent studies describing membrane remodeling by condensates in context of ESCRT biology (32,33). Together with its ability to tether membranes to spindle microtubules, these observations suggest that LEM2 condensation can generate membrane-remodeling forces that are spatially constrained by the structured LEM2-ESCRT-III scaffold during NE sealing.

**Figure 4:**
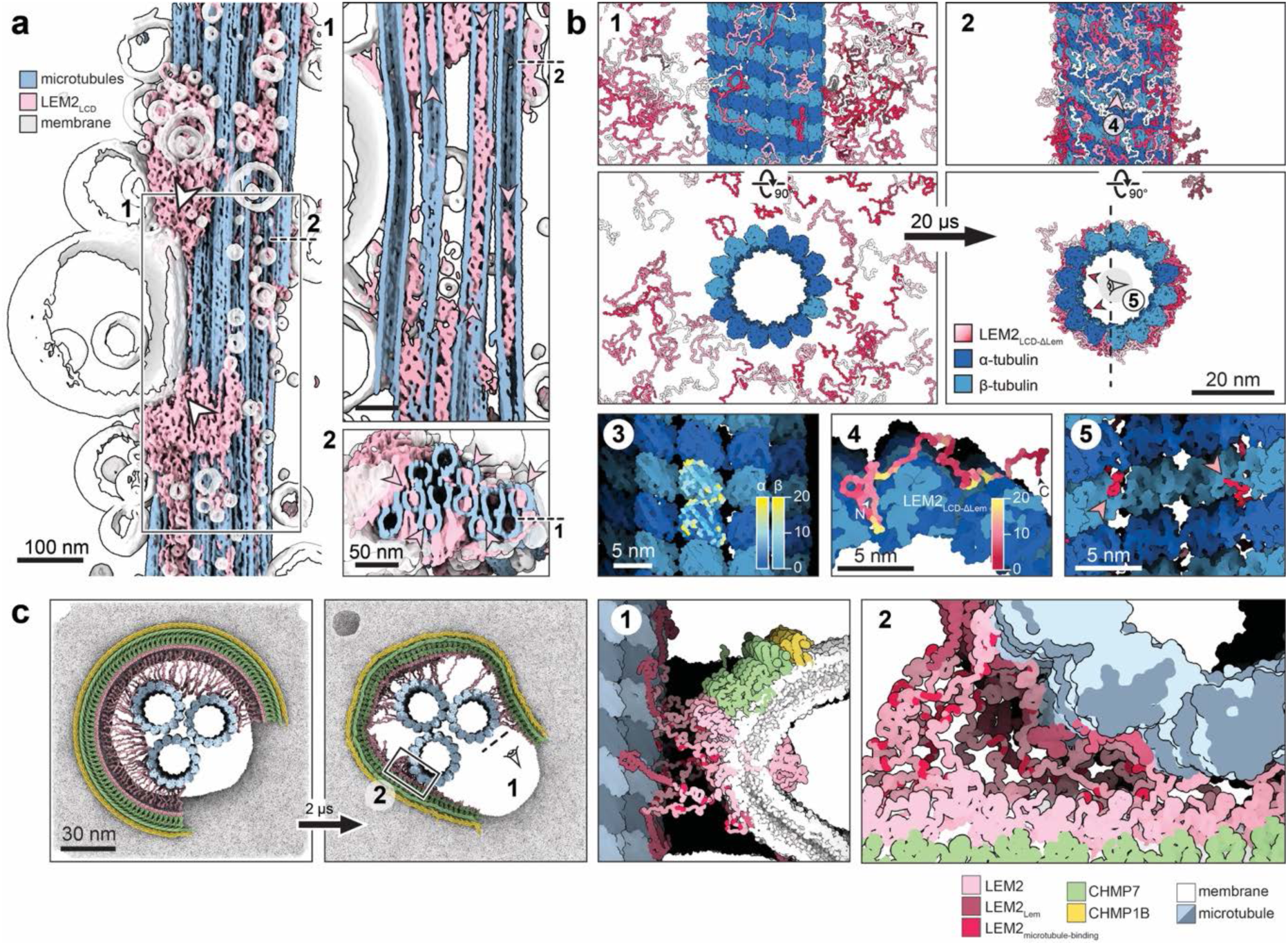
LEM2_LCD_ couples the membrane scaffold to microtubules. **a)** LEM2_LCD_-His_6_ condensates tether Ni-NTA vesicles to a microtubules bundle; tomographic segmentation. Longitudinal (1) and cross-sectional (2) cuts highlighting LEM2_LCD_-His_6_ density within the microtubule lumen. **b)** MD simulation: LEM2_LCD_ condenses on microtubules. Residue interactions are mapped in yellow on tubulin (3) and LEM2_LCD_ (4). LEM2_LCD-ΔLem_ molecules begin to enter through the lattice of protofilaments (5). **c)** Coarse-grained MD simulation of an open LEM2-CHMP7-CHMP1B filament on a membrane pore containing three microtubules. The filament tilts toward the pore neck while LEM2_LCD_ establishes extensive contacts with the microtubules, forming a dense protein meshwork that connects the membrane-bound scaffold to the spindle microtubules (1,2).

Unexpectedly, LEM2_LCD_-His_6_ also accumulated within the microtubule lumen (Fig. 4a-1,2, pink arrow heads, Supp. Fig. 12f-h), but only beneath external condensates, suggesting entry through the microtubule lattice rather than through microtubule ends. Coarse-grained MD simulations recapitulated rapid wetting of the microtubule surface and identified preferential interactions near gaps between protofilaments (Fig. 4b, Supp. Fig. 13). Five interaction patches were identified within LEM2_LCD_, three of those patches contained all four tryptophan residues of the LEM2_LCD_ sequence (Fig. 4b, Supp. Fig. 13b). During the simulations, individual LEM2_LCD-ΔLem_ molecules began migrating through the protofilament lattice toward the lumen (Fig. 4b, Supp. Fig. 13a), consistent with the experiments.

The preferential accumulation of LEM2_LCD_ within the lumen raises the possibility that microtubule lattice opening creates additional favorable interactions, potentially linking LEM2 wetting to subsequent spindle disassembly. Previous work established that ESCRT-III recruits the microtubule-severing ATPase spastin to nuclear sealing sites (5,34). Consistent with this idea, spastin accumulated within LEM2_LCD_ condensates assembled on microtubules *in vitro* (Supp. Fig. 14). Moreover, depletion of LEM2 delays spindle microtubule clearance during NE reformation in cells (8), supporting a role for the LEM2-organized sealing environment in coordinating subsequent microtubule disassembly.

We next asked how the LEM2-microtubule interaction influences pore geometry to form a diffusion barrier (8).

Coarse-grained MD simulations of LEM2-CHMP7-CHMP1B assembled on membrane pores surrounding three microtubules showed that LEM2_LCD_ rapidly engages microtubules, drawing the ESCRT-membrane scaffold towards them. This LEM2_LCD_-mediated tethering promoted the flat-to-tilted transition of the LEM2-CHMP7-CHMP1B co-polymer and markedly reduced the diffusion path between membrane and microtubules (Fig. 4c, Supp. Fig. 15). In contrast to simulations lacking microtubules, LEM2_LCD_ simultaneously restrained the filament ends at the microtubule surface, thereby stabilizing the entire tilted ESCRT scaffold. This resembled the tight interaction interface between NE membranes and microtubules observed in cells and *in vitro* (Supp. Fig. 1).

In summary, the intrinsically disordered LEM2_LCD_ mechanically couples the structured ESCRT scaffold to microtubules through wetting. The resulting forces promote scaffold remodeling while simultaneously narrowing the remaining diffusion path between membranes and microtubules. Once these microtubules are cleared, however, the physical constraint changes: the remaining membrane opening is no longer required to accommodate the spindle and can proceed toward terminal constriction. We therefore investigated how the LEM2-ESCRT-III machinery reorganizes after removal of the microtubule obstacle.

### LEM2 transitions from an ESCRT scaffold to a membrane-remodeling polymer

Terminal NE closure poses a distinct topologic challenge: ESCRT assemblies must transition from the flat membrane into the negatively curved inner surface of a constricting neck. High-resolution structures have predominantly captured ESCRT-III on positively curved membranes (21,35–37), and negative-curvature assemblies have required membrane deposition on preassembled protein tubes (24), leaving unresolved how the appropriate topology emerges during membrane remodeling itself.

In our membrane-bound LEM2_WH_-CHMP7-CHMP1B samples, however, we observed the appropriate topology in constricted membrane tubes, which were lined by protein spirals on their lumenal surface (Fig. 5a). To identify which of the three proteins generated these tubes, we systematically screened them in tomograms of different membrane reconstitution systems. To our surprise, neither of the ESCRT components, but LEM2_WH_ alone, remodeled membranes into such tubes (Fig. 5b, Supp. Fig. 16). Subtomogram averaging yielded a 8.3-Å reconstruction of LEM2_WH_ tubes (Methods) and enabled rigid-body fitting of the LEM2_WH_ domain (Fig. 5c, Supp. Fig. 17). This revealed that the tubes were predominantly composed of two-start helices formed by triple-stranded assemblies in either parallel or antiparallel orientations (Supp. Fig. 17).

**Figure 5:**
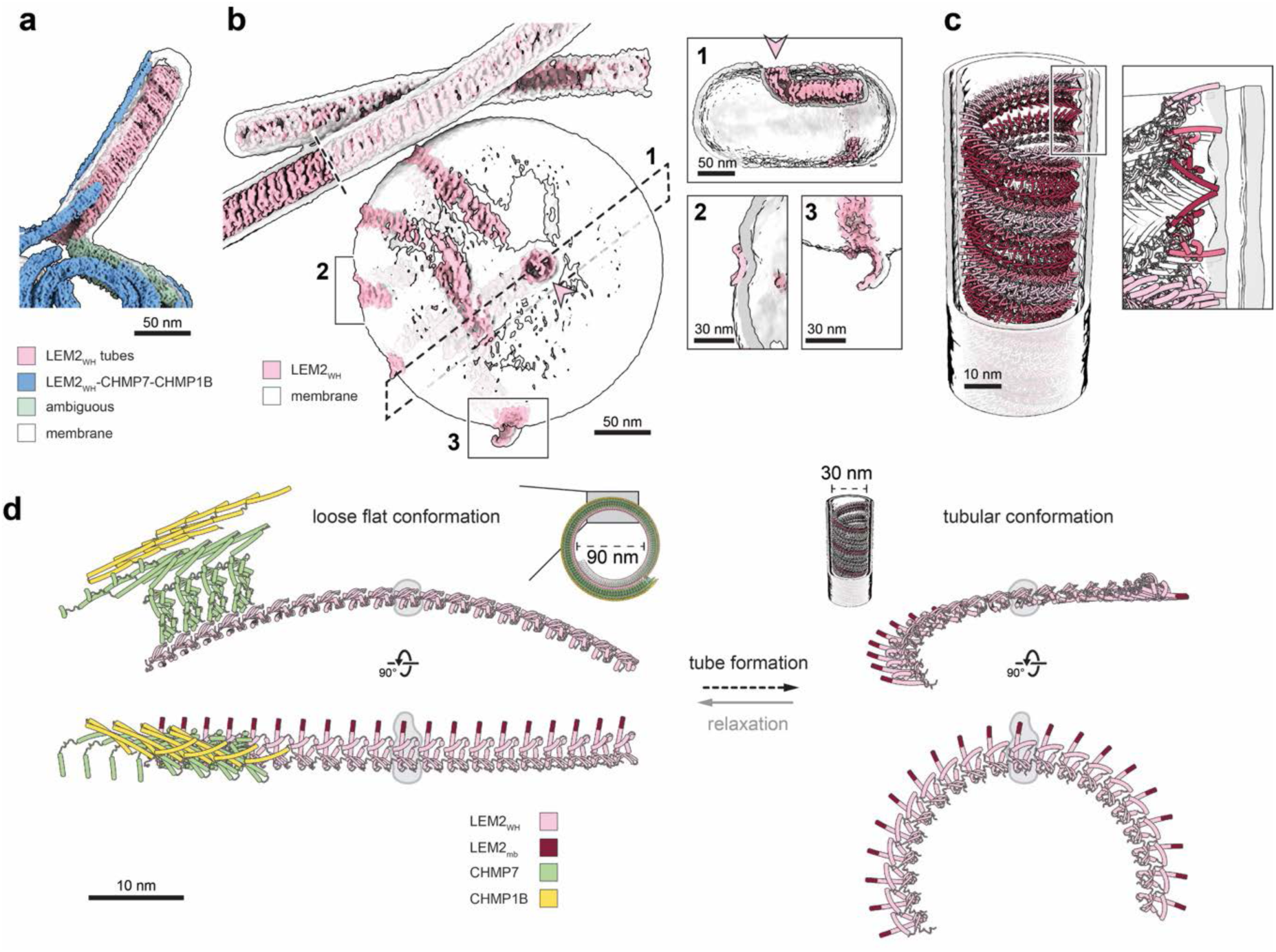
LEM2_WH_ polymerization generates tightly constricted, negatively curved membrane tubes. **a)** Reconstitution of LEM2_WH_, CHMP7 and CHMP1B on vesicles produces tightly constricted membrane tubes with negative curvature; tomographic segmentation. **b)** Reconstitution of LEM2_WH_ alone on vesicles demonstrates that LEM2_WH_ is sufficient to form highly curved membrane-associated filaments and tubes; tomographic segmentation, (1) LEM2_WH_ assemblies generate openings in vesicles. (2) At the vesicle periphery, LEM2_WH_ forms curved filaments whose cross-sections resemble incomplete tubular assemblies. (3) Peripheral LEM2_WH_ filaments deform and pull membrane from vesicles as they transition toward a tubular architecture. **c)** Subtomogram average of antiparallel LEM2_WH_ tubes formed in reconstitutions containing LEM2_WH_, CHMP7 and CHMP1B, with rigid-body fitting of the structural model. **d)** Comparison of LEM2_WH_ strand conformation in filaments with CHMP7 and CHMP1B (left) versus alone in tubes (right).

Our LEM2_WH_-CHMP7-CHMP1B structure identified LEM2 as an iso-stoichiometric component of the ESCRT-III scaffold itself (Fig. 2), and we demonstrated that remodeling of the ESCRT-III filament simultaneously remodels the LEM2 polymer (Fig. 3e, 4c). This raises the possibility that beyond its recruiting and scaffolding role, LEM2 is involved in late-stage membrane remodeling and neck constriction in NE formation. Consistent with this idea, others have shown that Vps4-mediated release of LEM2 from CHMP7 is required for re-establishing nucleocytoplasmic compartmentalization (10,15). With our data, this suggests a model where ESCRT-III stabilizes pore geometries to prime an intrinsic membrane-remodeling state of LEM2 during terminal pore constriction. To address this, we compared LEM2 organization between filaments formed by LEM2_WH_-CHMP7-CHMP1B and LEM2_WH_ alone. This revealed that a transition from large-curvature to tube conformations requires only minor rearrangements of lateral contacts and neighboring subunit angles, which lead to a rotation of filament curvature relative to the subunit axis (Fig. 5d). Cohesively, this conformational transition corresponds to the flat-to-tilted conformational change LEM2 undergoes in the MD simulations (Fig. 3e). Hence, the experimentally determined organization of helical LEM2_WH_ filaments supports the flat-to-tilted transition that is suggested by the MD simulations. Further support comes from the rare late-stage intermediate with detached microtubules yet open membrane pore on locally condensed chromatin (Supp. Fig. 2). Here, the membrane pore showed a constricted state corresponding to the tube diameter of helical assemblies formed by LEM2_WH_ alone (Supp. Fig. 16d). Together, this suggests that the LEM2-ESCRT-III scaffold directly precedes formation of a tubular LEM2 polymer during membrane pore maturation. We therefore propose a sequential mechanism for NE sealing (Fig. 6): LEM2 is first recruited to spindle microtubules through its LCD, to co-polymerize with CHMP7 and produce large curvature membrane scaffolds (Fig. 6a,b). Recruitment of downstream ESCRT-III proteins such as CHMP1B promotes tilting of the scaffold as it adapts to the shrinking membrane neck (Fig. 6c,d). As the filament is being remodeled, LEM2 transitions into tightly curved membrane-remodeling helices that drive terminal pore constriction (Fig. 6e).

**Figure 6:**
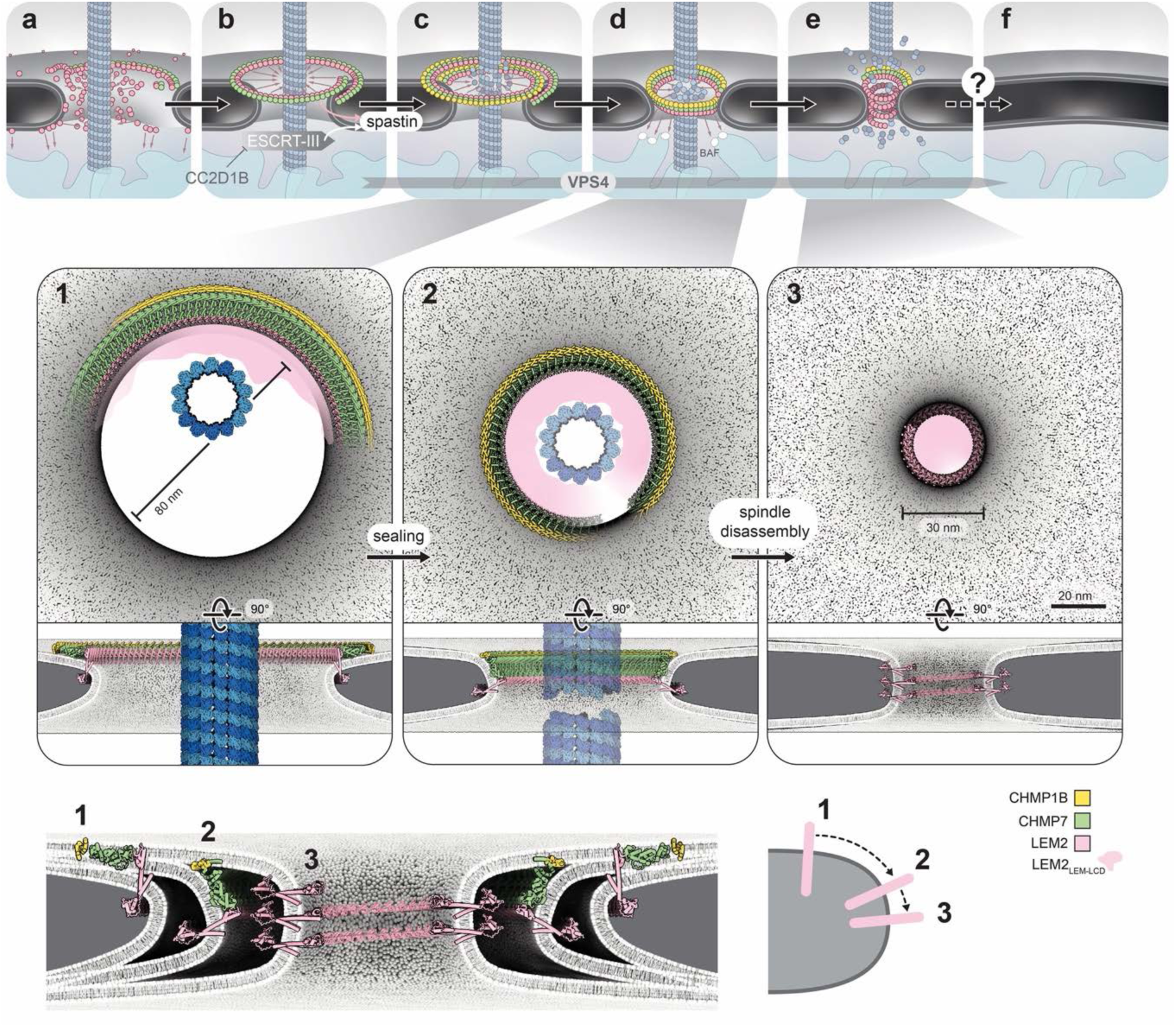
Model for stepwise nuclear-envelope closure orchestrated by LEM2–ESCRT-III. **a)** LEM2 condenses around spindle microtubules through interactions mediated by LEM2_LCD_. **b)** CHMP7 binding nucleates a membrane-associated LEM2–ESCRT-III scaffold around the pore. **c)** CHMP1B incorporation stabilizes the filament and establishes a membrane-bound scaffold spanning the pore; structural model derived from cryo-EM, subtomogram averaging and MD simulations (1). **d)** The scaffold transitions from a flat to a tilted configuration and migrates toward the pore neck, adapting its geometry to the catenoid membrane topology; structural model derived from MD simulations (2). **e)** Following spindle-microtubule clearance, LEM2WH polymerization drives formation of tightly curved membrane-associated tubes and further pore constriction; structural model derived from cryo-EM and subtomogram averaging (3). **f)** Final membrane scission and pore closure proceed by a mechanism that remains unresolved.

## Discussion

To re-establish nucleocytoplasmic compartmentalization at mitotic exit, the reforming NE must seal around spindle microtubules that mechanically prevent complete membrane closure. Here, we show how this is coordinated by structured polymers that guide membrane pore constriction and intrinsically disordered protein domains that seal the diffusion path. Specifically, the transmembrane receptor LEM2 co-polymerizes with CHMP7 to form a membrane-bound scaffold that stabilizes NE openings and recruits downstream ESCRT-III proteins, while positioning its LCD within the pore. There, LEM2 condenses around spindle microtubules, mechanically couples the membrane scaffold to its obstacle and narrows the remaining diffusion path. Following microtubule clearance, LEM2 can transition into tightly curved membrane-remodeling polymers. Thus, rather than acting simply as an ESCRT receptor, LEM2 organizes successive molecular states that connect spindle enclosure, transient sealing and terminal membrane constriction.

This mechanism suggests a broader principle for ESCRT-mediated membrane remodeling. ESCRT adaptors are generally considered to specify where ESCRT-III assembles, whereas membrane geometry is attributed primarily to the ESCRT-III polymers themselves. Our structures instead show that the receptor can become an integral component of the polymer and directly determine its architecture. Co-polymerization with CHMP7 maintains LEM2 in a large-curvature scaffold compatible with pores containing microtubules, whereas changes in filament composition progressively constrain the scaffold and favor its transition towards the membrane neck. LEM2 alone can subsequently form tightly curved polymers lining negatively curved membrane tubes. Membrane receptors may therefore encode not only the location but also the topology and progression of ESCRT-mediated membrane remodeling. How pore closure is ultimately completed, including membrane scission and fusion, remains to be determined. Intriguingly, LEM2_LCD_ induces membrane blebbing and cargo-containing intraluminal vesicle formation in our reconstitutions (Supp. Fig. 12d,e), raising the possibility that the disordered region contributes directly to late membrane-remodeling events. Condensates can themselves drive membrane deformation and scission, suggesting that LEM2 condensation may contribute to terminal closure (33). Our observation of LEM2 density within the microtubule lumen further suggests that condensation along the microtubule surface can promote access through lattice defects or openings. Together with our previous finding that LEM2 condensation generates substantial forces on DNA, these observations suggest a common physical mechanism in which multivalent electrostatic interactions allow the LEM2_LCD_ to condense at DNA, microtubule and membrane interfaces and convert condensation into mechanical work. Depending on the geometry of the substrate, this may reinforce DNA-membrane contacts, couple membranes to microtubules, or drive membrane deformation (8,38).

This regulation also reveals a double-edged nature of LEM2-CHMP7 activity. Premature formation of tightly constricted LEM2 polymers would be incompatible with spindle enclosure, suggesting that ESCRT assembly initially restrains an intrinsic membrane-remodeling activity of LEM2 and channels it through geometries appropriate for each stage of closure. Such control may be particularly important because excessive LEM2-CHMP7 activity is itself damaging. In micronuclei, unrestrained CHMP7-LEM2 accumulation is associated with aberrant membrane deformation, chromosome fragmentation and ultimately chromothripsis, whereas defective NE assembly is a major determinant of micronuclear instability (39–41). Our structures provide a possible molecular basis for this balance: the conformational plasticity that enables productive remodeling during NE closure could become destructive when spatially or temporally misregulated. Recent work further shows that LEM2, CHMP7 and CHMP1B assemble around mis-segregated chromatin into DNA-membrane bridges with geometries distinct from those described here (42). The involvement of CHMP1B in both assemblies, yet in distinct experimentally determined configurations, further demonstrates the remarkable conformational range accessible to ESCRT proteins and suggests that it is shaped by biological context.

A second principle emerging from our work is the cooperation between ordered and disordered molecular assemblies. The LEM2-ESCRT scaffold generates a circumferential array of LCDs that condense around spindle microtubules and fill the irregular space that cannot yet be eliminated by membrane fusion. A structured polymer therefore defines the geometry of the reaction, whereas a dynamic LCD meshwork establishes a functional diffusion barrier before membrane continuity is restored. This arrangement is conceptually reminiscent of the NPC, where a rigid scaffold organizes a dense phase of intrinsically disordered FG-repeat nucleoporins to establish selective compartmentalization. More broadly, our findings illustrate how cells can combine polymerization, condensation and membrane remodeling to progressively transform a topologically loosely defined opening into a sealed organelle boundary.

## Acknowledgement

We thank Adam Frost and Martin Beck for their generous support and feedback throughout the course of this project. From the MPI-CBG, we thank Petra Kiesel and Swantje Lenz, Oscar Gonzales from the IT department, Tobias Fürstenhaupt and Matthias Poge from the EM facility for their technical support. Iterative system optimization and simulation data processing were supported by the Max Planck Computing and Data Facility (MPCDF).

## Funding

This research was supported by the Max Planck Society (AvA, GH), European Research Council, Organelloids 101117619 (AvA), the German Research Foundation SPP2191 (AvA) and the “Hessen Horizon Marie Skłodowska-Curie-Stipendium” program (KPR). We performed large-scale simulations on the national supercomputer HPE Apollo Hawk at the High Performance Computing Center Stuttgart (HLRS) under grant GCS-MDhNPC/ACID 44255.

---

**Supp. Figure 1:**
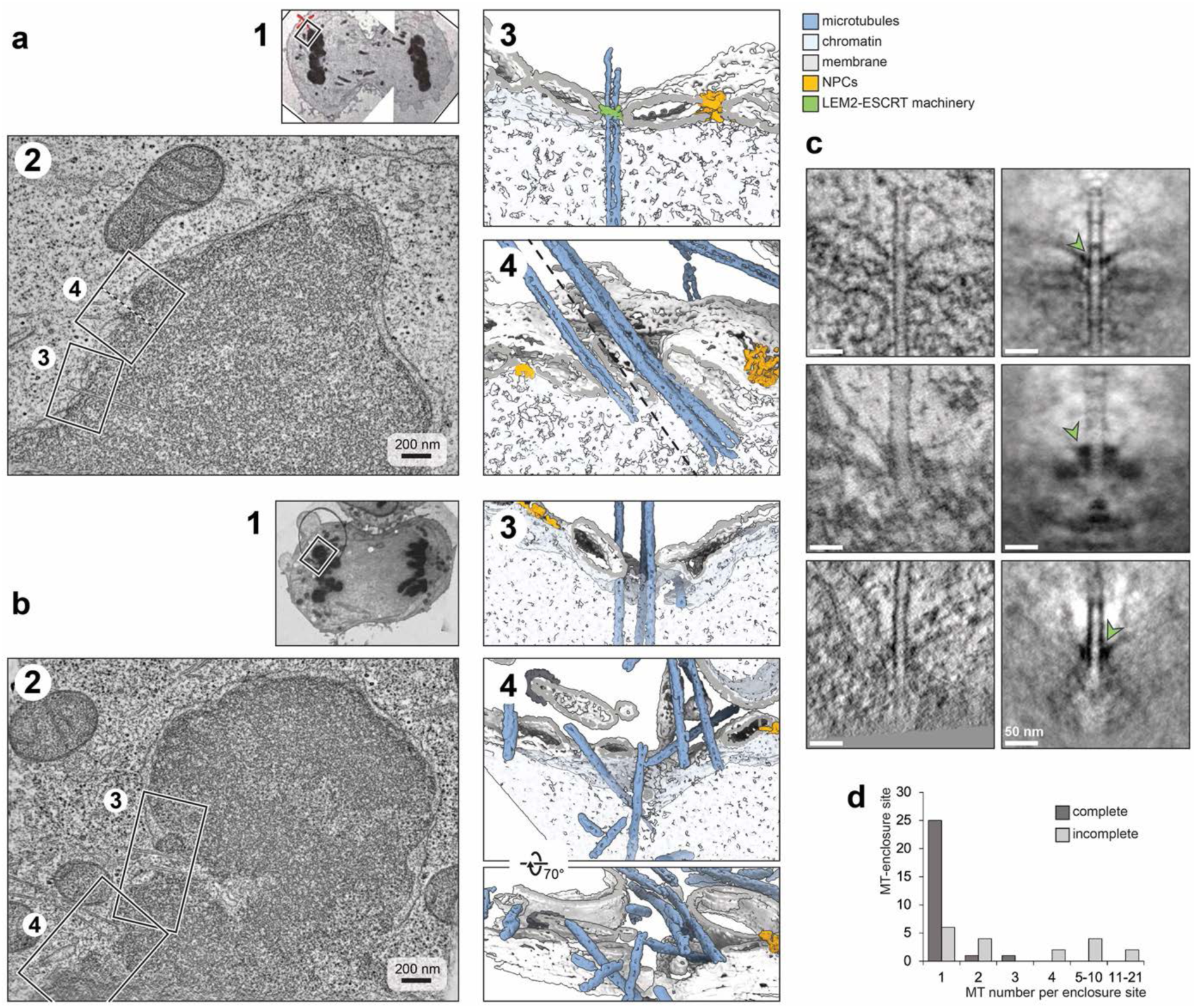
Topology of spindle MT entry sites through the NE during anaphase. **a,b)** Tomograms of microtubule entry events in high-pressure-frozen HeLa cells at 7.7 min **(a)** and 6.2 min **(b)** after anaphase onset. 1) Overview of the HeLa cell. 2) Tomographic slice of the region indicated in 1. 3,4) Segmentations of microtubule-enclosure sites at the NE, as indicated in 2. a3) A single microtubule is enclosed by the NE at a near-perpendicular orientation. A ring-like density associated with the putative LEM2–ESCRT machinery is highlighted in green (the same example is shown in c, left). **a4)** Close-up view of two microtubule-enclosure sites with one (left) and two (right) microtubules passing through individual NE openings. Note the elongated shape of the NE around the microtubules. Left and right parts of the panel are shown at different clipping heights. **b3)** Three parallel microtubules extend through one NE opening, while another single microtubule extends through a separate opening. **b4)** Multiple microtubules remain at different angles within NE openings of varying size. **c)** Single-microtubule enclosure events in which the microtubules are oriented approximately perpendicular to the NE (top), and their rotational averages (bottom). The microtubule is located centrally and perpendicular to the NE plane; chromatin is in the lower half and cytoplasm in the upper half of the panels, separated by the NE. Green arrowheads indicate density associated with the putative LEM2-ESCRT machinery. **d)** Microtubule-enclosure events were counted and plotted against the number of microtubules per event. Most microtubule-enclosure events contained a single microtubule. However, larger bundles of 5-21 or more microtubules were also observed and likely represent earlier-stage intermediates. Dark grey: the microtubule-enclosure site was fully contained within the tomogram, allowing all microtubules in the respective bundle to be counted. Light grey: the microtubule bundle was only partially contained within the tomogram and therefore likely comprised more microtubules than indicated. Events were counted from 20 tomograms acquired from four cells between 6.1 and 9.3 min after anaphase onset.

**Supp. Figure 2:**
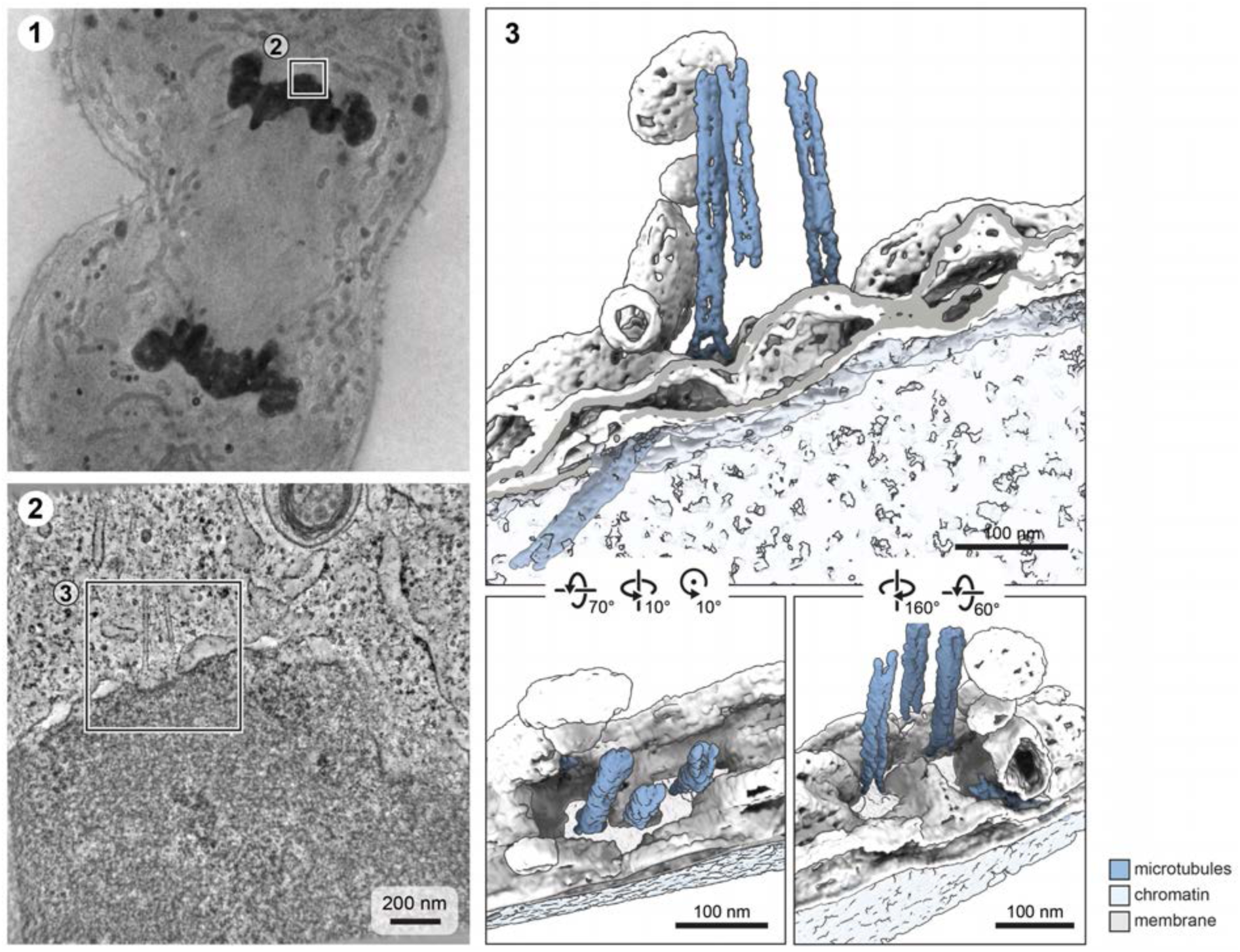
Terminal geometry of a double-membrane pore following spindle-microtubule detachment. **a)** Tomogram of a high-pressure-frozen HeLa cell at 8.4 min after anaphase onset, showing a double-membrane pore shortly after detachment of spindle microtubules. 1) Overview. 2) Tomographic slice. 3) Segmentation shown from different viewing angles. Three microtubules remain positioned in the cytoplasm above a shared NE pore, whose shape still reflects the geometry of the preceding MT bundle. A fourth MT traverses the chromatin and passes through the NE within a separate pore.

**Supp. Figure 3:**
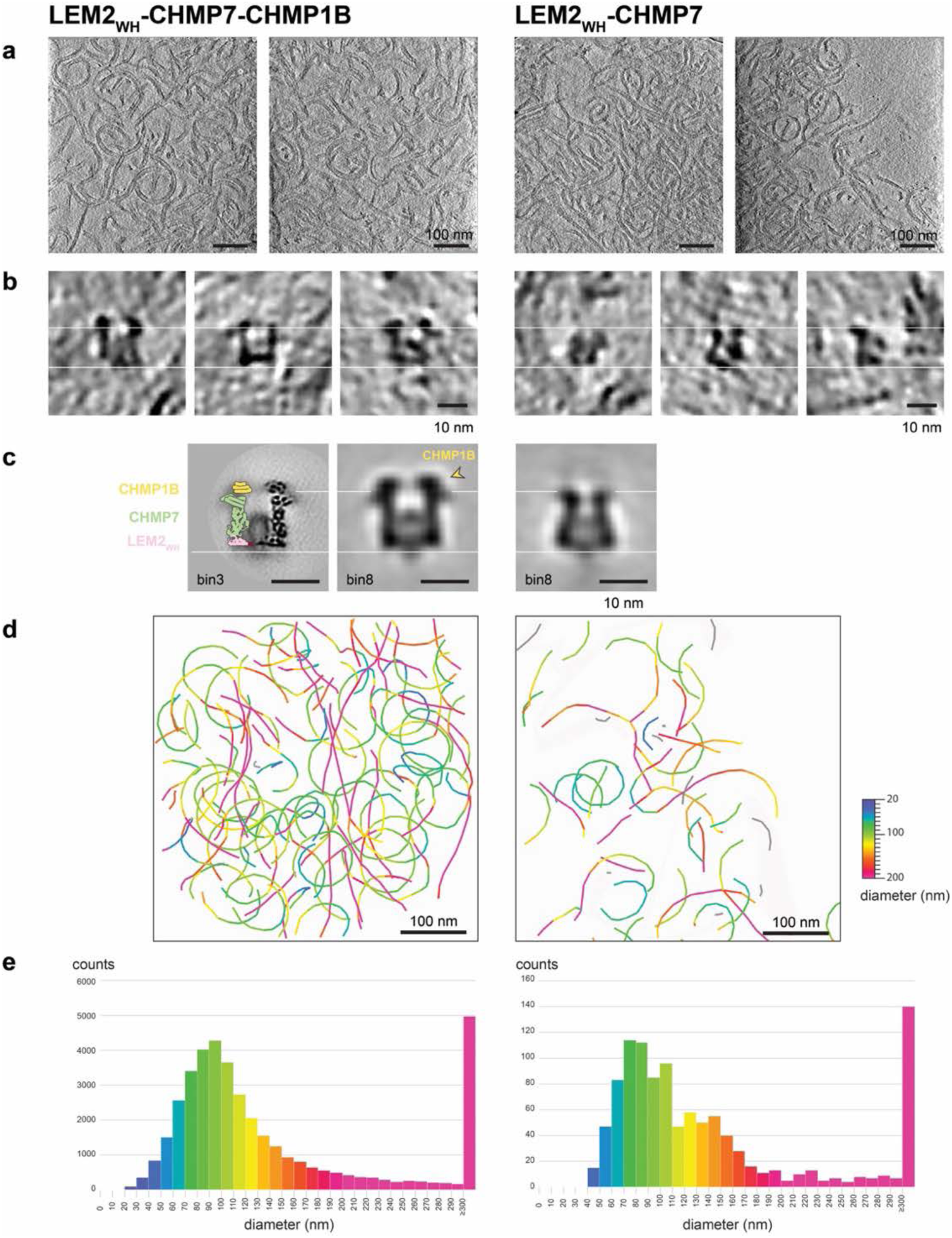
LEM2_WH_-CHMP7-CHMP1B and LEM2_WH_-CHMP7 filaments have similar architectures and preferred curvatures. **a)** Tomographic slices of LEM2WH–CHMP7–CHMP1B (left) and LEM2WH–CHMP7 filaments (right) assembled in the presence of solubilized lipids. Slice thickness: 10 px. **b)** Tomographic slices showing cross sections of LEM2WH–CHMP7–CHMP1B filaments (left) and LEM2_WH_–CHMP7 filaments (right). White lines serve as guides for comparing filament architecture and highlight additional density in the CHMP1B-containing filaments above the upper line. **c)** Subtomogram averages show that LEM2_WH_-CHMP7 assembles into filaments with an overall architecture similar to that of LEM2_WH_-CHMP7–CHMP1B. In the absence of CHMP1B, the two symmetric filament strands are slightly more tilted relative to each other. For direct comparison, bin8 averages were generated from 5,000 particles per condition. Slice thicknesses of 10 px and 5 px are shown for the bin8 and bin3 averages, respectively. **d)** 3D traces of filaments in a tomogram, color-coded according to local filament diameter/curvature. Straight filament segments (pink) predominantly correspond to filaments located at the air– water interface, where the preferred curvature is reduced. **e)** Histograms of local filament diameters measured using a 100-px window in bin8 tomograms. Both filament compositions display similar preferred diameters (LEM2_WH_-CHMP7-CHMP1B: 80-100 nm; LEM2_WH_-CHMP7: 70-90 nm). The LEM2_WH_–CHMP7 distribution is broader and noisier, indicating that filaments lacking CHMP1B sample a wider range of curvatures. Most LEM2_WH_–CHMP7– CHMP1B data were acquired on a Titan Krios, whereas LEM2_WH_-CHMP7 data were acquired on a Titan Halo. To enable direct comparison, the data shown in a, b and the bin8 averages in c were acquired on a Titan Halo for both conditions.

**Supp. Figure 4:**
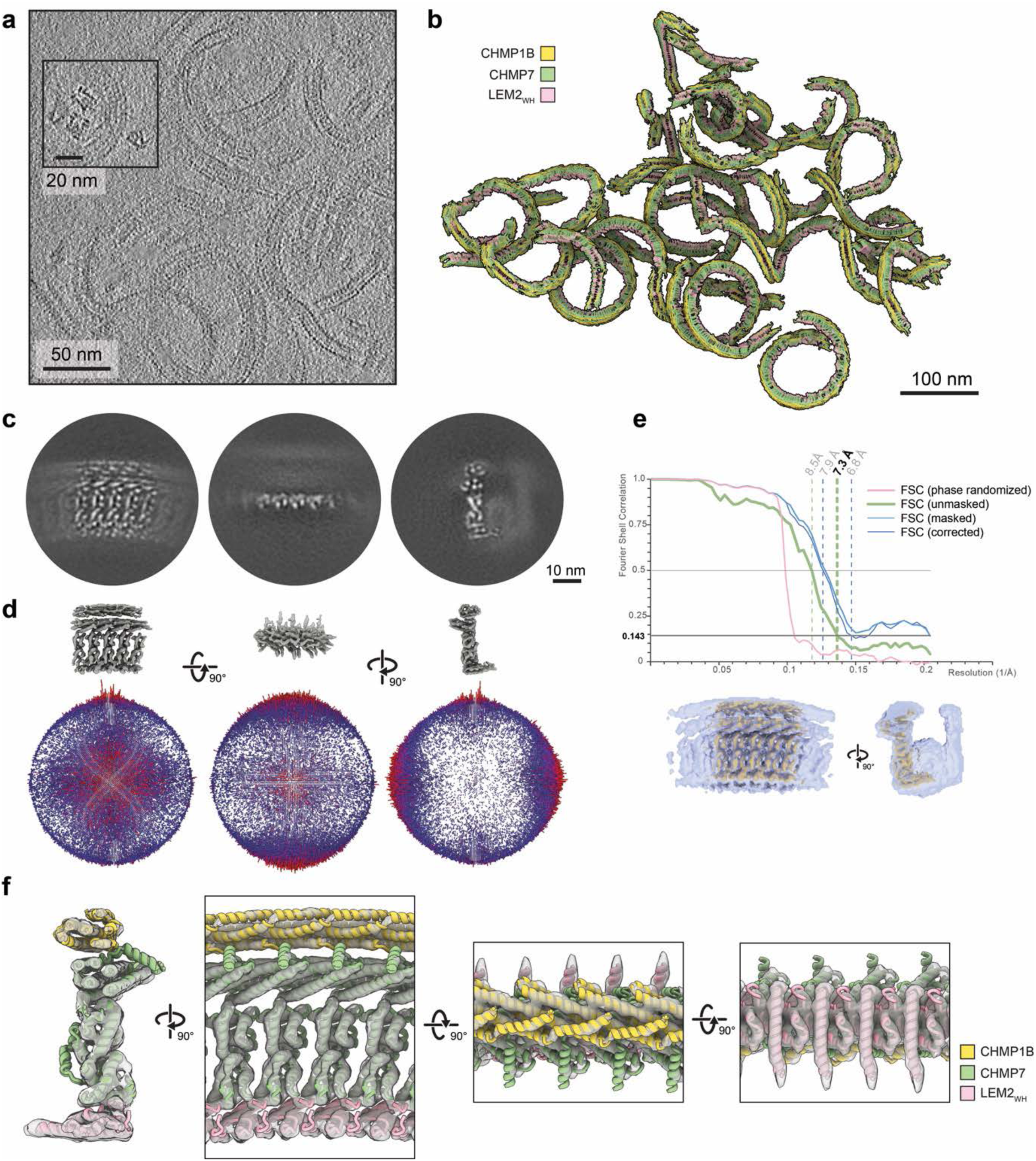
Tomograms and subtomogram averaging of LEM2_WH_-CHMP7-CHMP1B filaments in presence of solubilized lipids. **a)** Tomographic slice, showing the repetitive and curved architecture of the filaments; the inset highlights representative cross-sectional views. **b)** The subtomogram average was mapped back onto the particle positions in the tomogram, revealing the overall filament architecture, with a preferred curvature but variation in twist and pitch. **c)** Central slices through the final subtomogram average of one filament strand. **d)** Volume (top) and angular distribution of particles (bottom). **e)** Fourier shell correlation (FSC) curves, reporting a resolution of 7.3 Å (top), and the mask (blue) that was used for the analysis (bottom). f) Symmetrized composite density used for modeling, with the atomic model fitted into the density.

**Supp. Figure 5:**
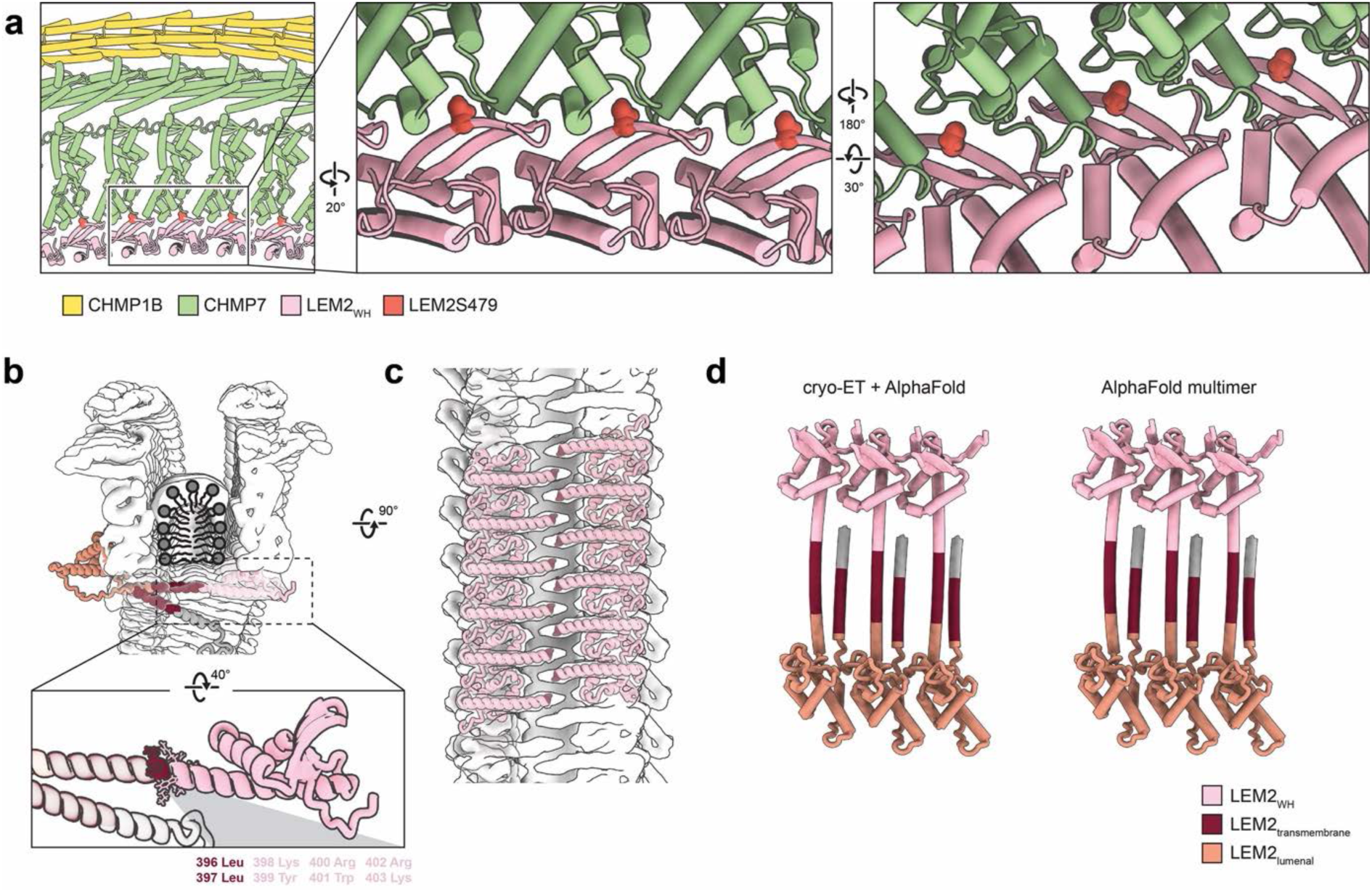
Interactions of LEM2. **a)** The progeria-associated mutation LEM2S479F is located within the interface between LEM2 and CHMP7. **b)** Placement of full-length LEM2 into the EM density of the anti-parallel LEM2_WH_-CHMP7-CHMP1B filament results in a clash between LEM2_TM_ and the opposite anti-parallel filament strand. The N terminus of the LEM2_WH_ construct retains eight residues from the adjacent transmembrane region: two leucine residues predicted to partition into the hydrophobic core of the lipid bilayer and six residues predicted to interact with phospholipid head groups. The bicelle-like density at the center of the C2-symmetric filament is indicated by schematic phospholipids (grey). **c)** The N-termini of LEM2_WH_ form a zipper with the opposite anti-parallel filament strand, in which the hydrophobic residues interact. **d)** Polymeric assembly of full-length LEM2. Left: Lateral assembly as indicated by the LEM2_WH_ cryo-ET density in filaments with CHMP7 and CHMP1B, and extended to their full-length structure as obtained from monomeric AlphaFold predictions. Right: AlphaFold Multimer prediction of three full-length LEM2 molecules, confirming the lateral placement suggested by the LEM2_WH_-ESCRT filament. Assignment of the core transmembrane domains of LEM2 (dark red) as predicted by DeepTMHMM-1.0 (43).

**Supp. Figure 6:**
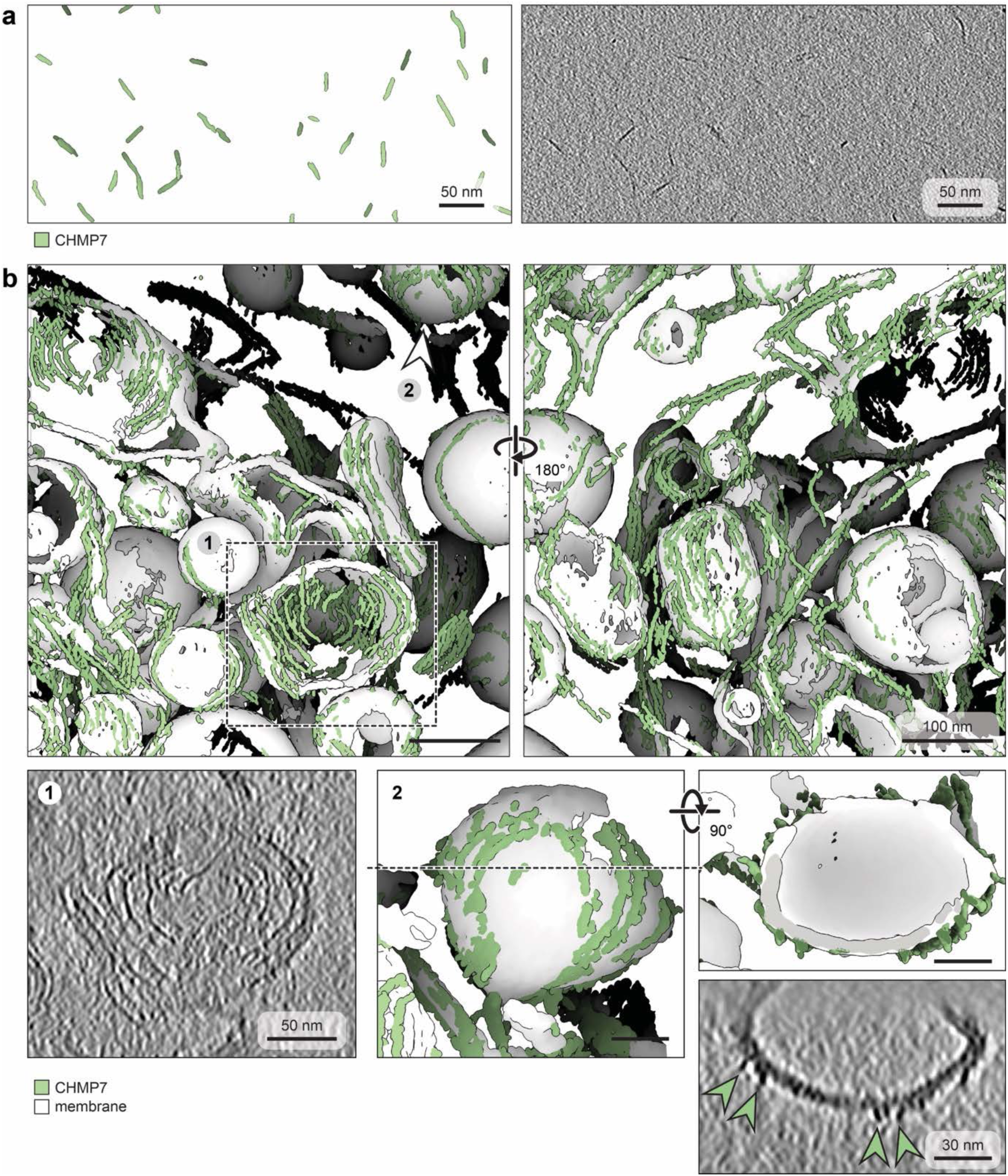
CHMP7 polymerization. **a)** CHMP7 polymerizes into short filamentous assemblies in the presence of solubilized lipids. **b)** On vesicles, CHMP7 polymerizes into extended, predominantly flat spiral filaments with a thin single-stranded appearance. The filaments locally deform and flatten the vesicle membrane. Representative tomographic segmentations and slices are shown; arrowheads indicate membrane-associated CHMP7 filaments.

**Supp. Figure 7:**
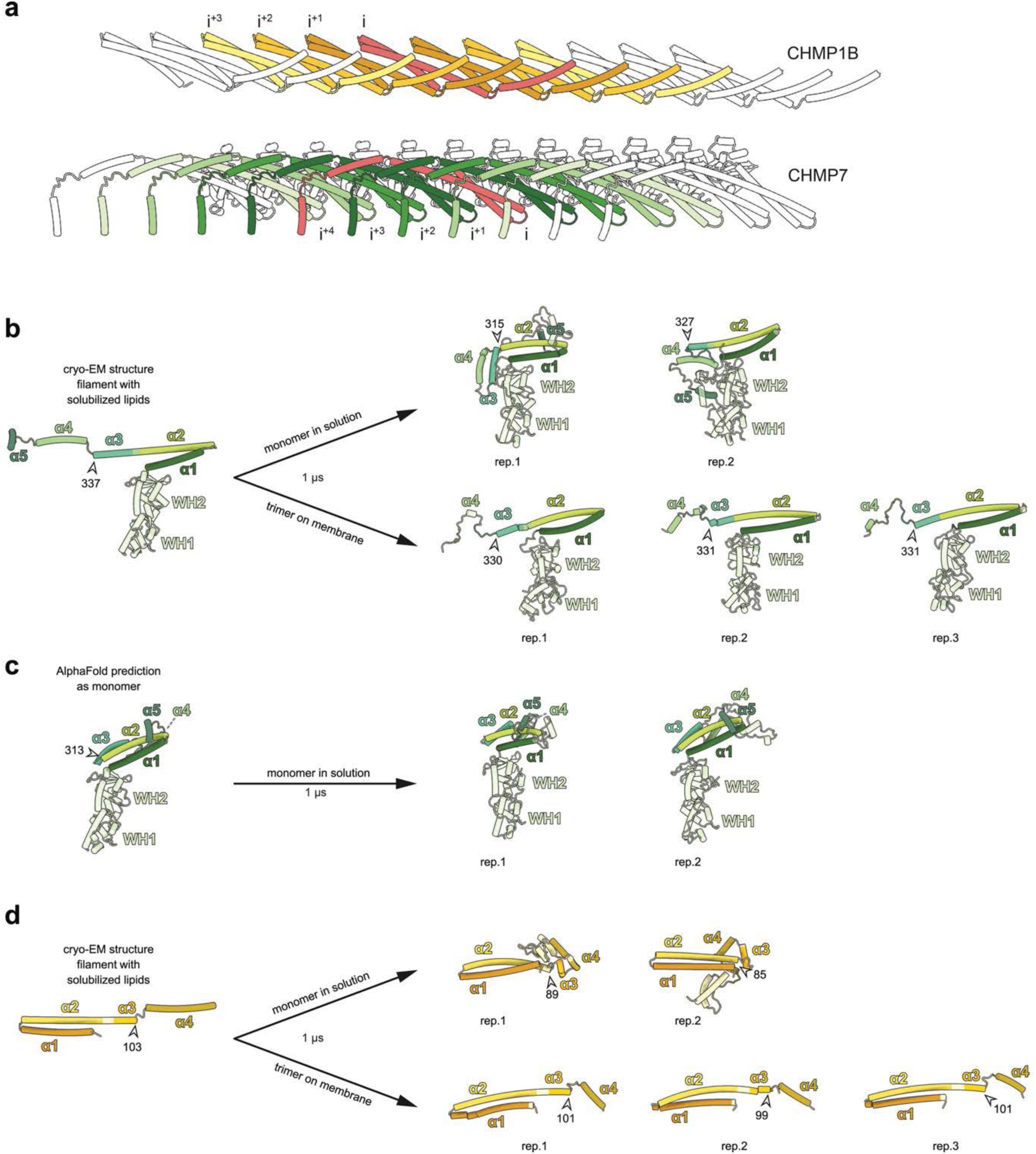
Polymer contacts stabilize the open conformations of CHMP7 and CHMP1B ESCRT-III domains. **a)** Within a filament of LEM2_WH_-CHMP7-CHMP1B, CHMP1B has lateral contacts with four neighboring subunits (top), and CHMP7 with eight neighboring subunits (bottom). Note that helix a5 of CHMP1B is not resolved in our structure and could provide contacts to further neighbors. **b-d)** All-atom simulations of ESCRT-III subunits over a time course of 1 µs. **b)** Simulation of the open CHMP7 conformation as derived from the cryo-EM structure results in breakage of the extended helix a2/a3 when put as monomer in solution, but remains more stable within a trimer of LEM2-CHMP7-CHMP1B on flat membrane. **c)** Simulation of the closed CHMP7 conformation as derived from AlphaFold remains stable in solution. **d)** Simulation of the open CHMP1B conformation as derived from the cryo-EM structure results in breakage of the extended helix a2/a3 when put as monomer in solution, but remains more stable within a trimer of LEM2-CHMP7-CHMP1B on flat membrane. For trimer simulations, the center subunit only is shown, as it experienced most stabilization through its neighbors.

**Supp. Figure 8:**
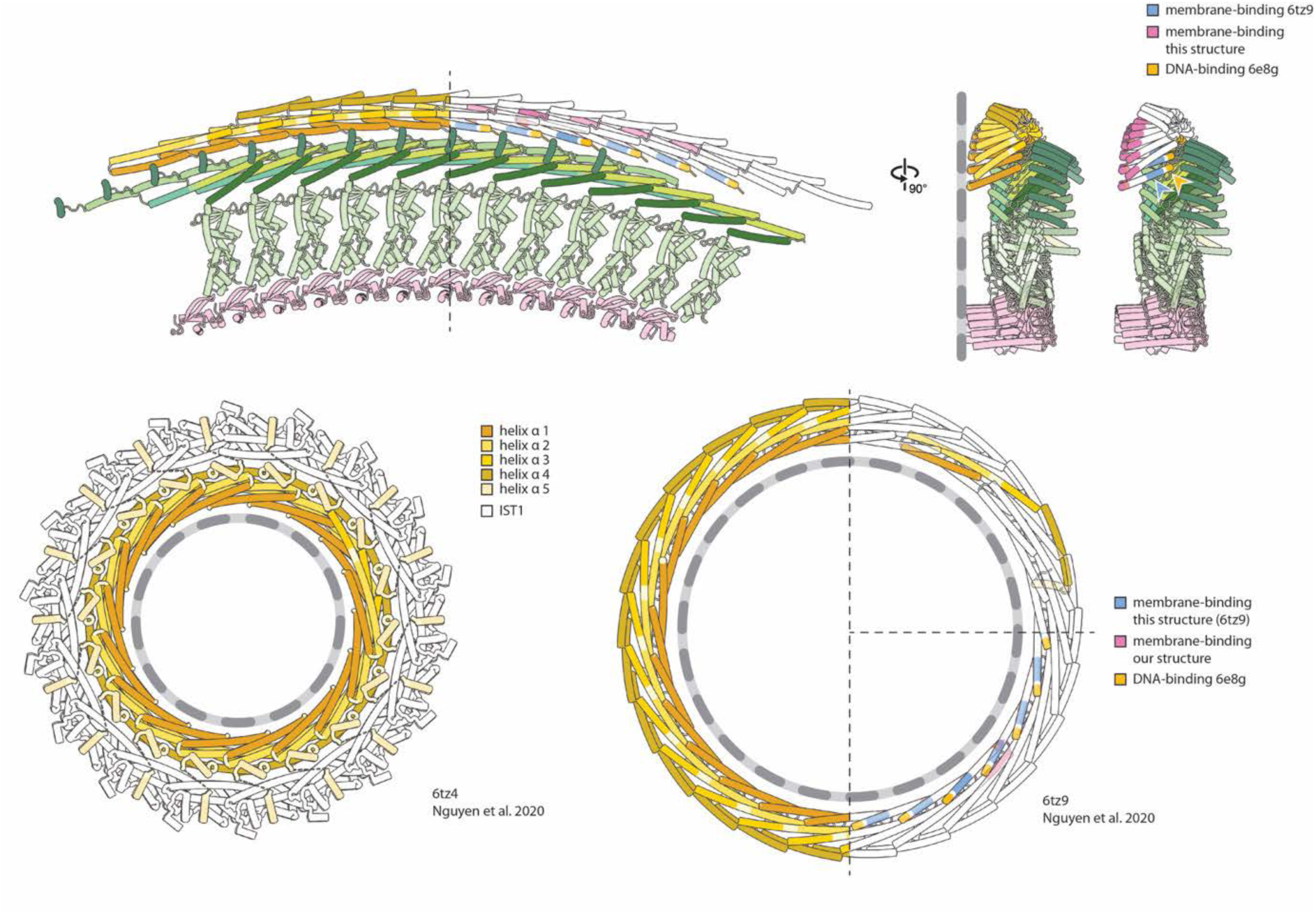
Comparison of CHMP1B membrane binding sites in different contexts. **a)** In a filament of LEM2-CHMP7-CHMP1B, CHMP1B has a membrane binding site at the loop between helix a1 and a2 (magenta). **b)** CHMP1B with IST1 (left) or without (right) forms membrane tubes with positive curvature (27) and has a membrane interaction site in helix a1 (blue). The binding site at the loop between helix a1 and a2 is buried in the interface to neighboring subunits. Vice versa, the binding site in helix a1 is buried in the interface to CHMP7 in our structure. This indicates that the CHMP1B filament has to undergo a conformational change to switch from one membrane binding face to the other.

**Supp. Figure 9:**
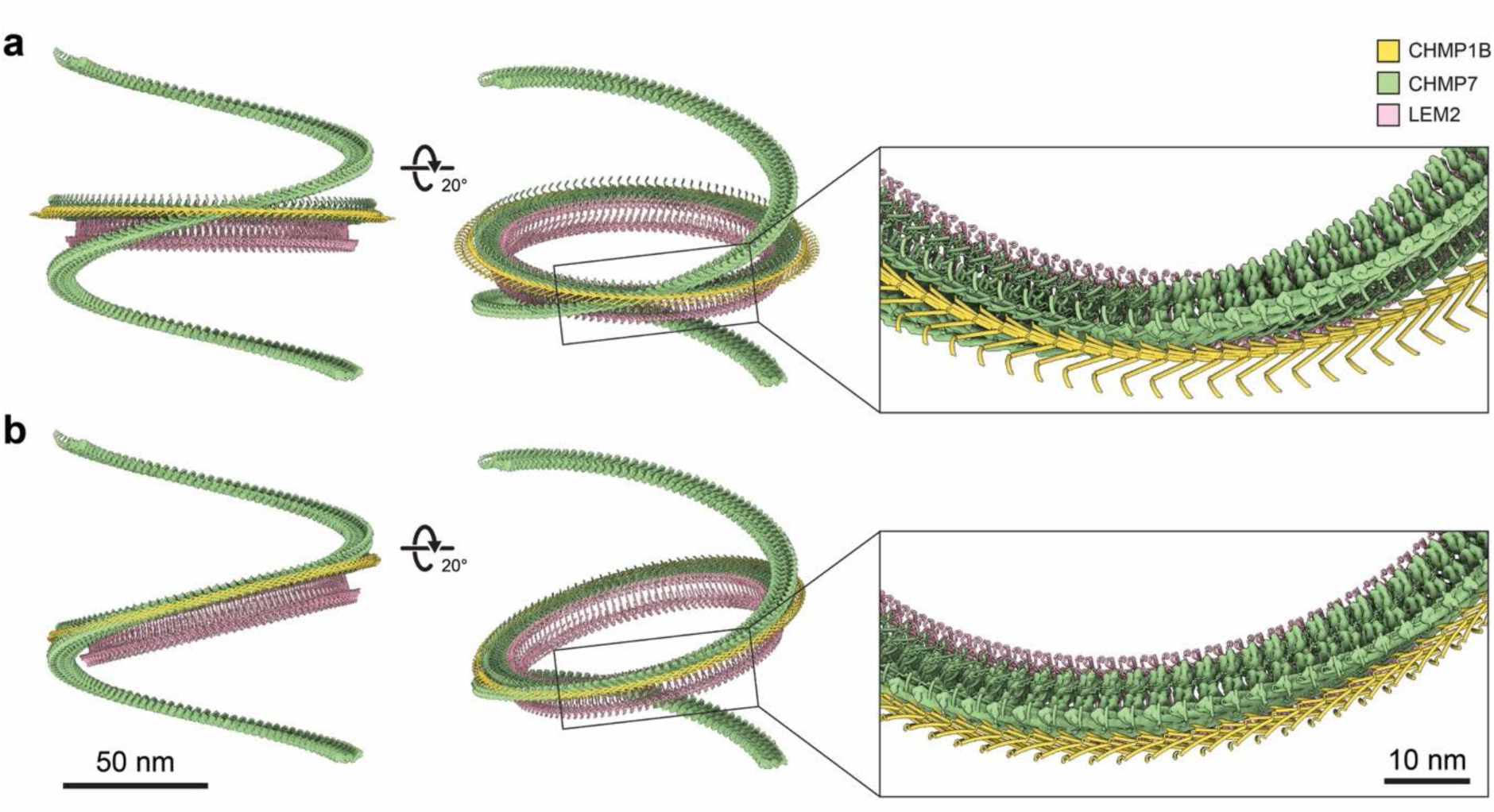
Adjusting the helical filament to accommodate a flat membrane geometry. **a)** Planar symmetrization around the helical axis resulted in substantial shifts of individual subunits relative to the original cryo-ET structure and poor conservation of lateral interaction. **b)** Instead, planar symmetrization was performed around an axis perpendicular to a peripheral filament plane. This resulted in only minimal shifts between subunits while preserving the lateral interactions. For simplicity, only a single-stranded CHMP7 density is shown to indicate the original helical architecture of the cryo-ET structure.

**Supp. Figure 10:**
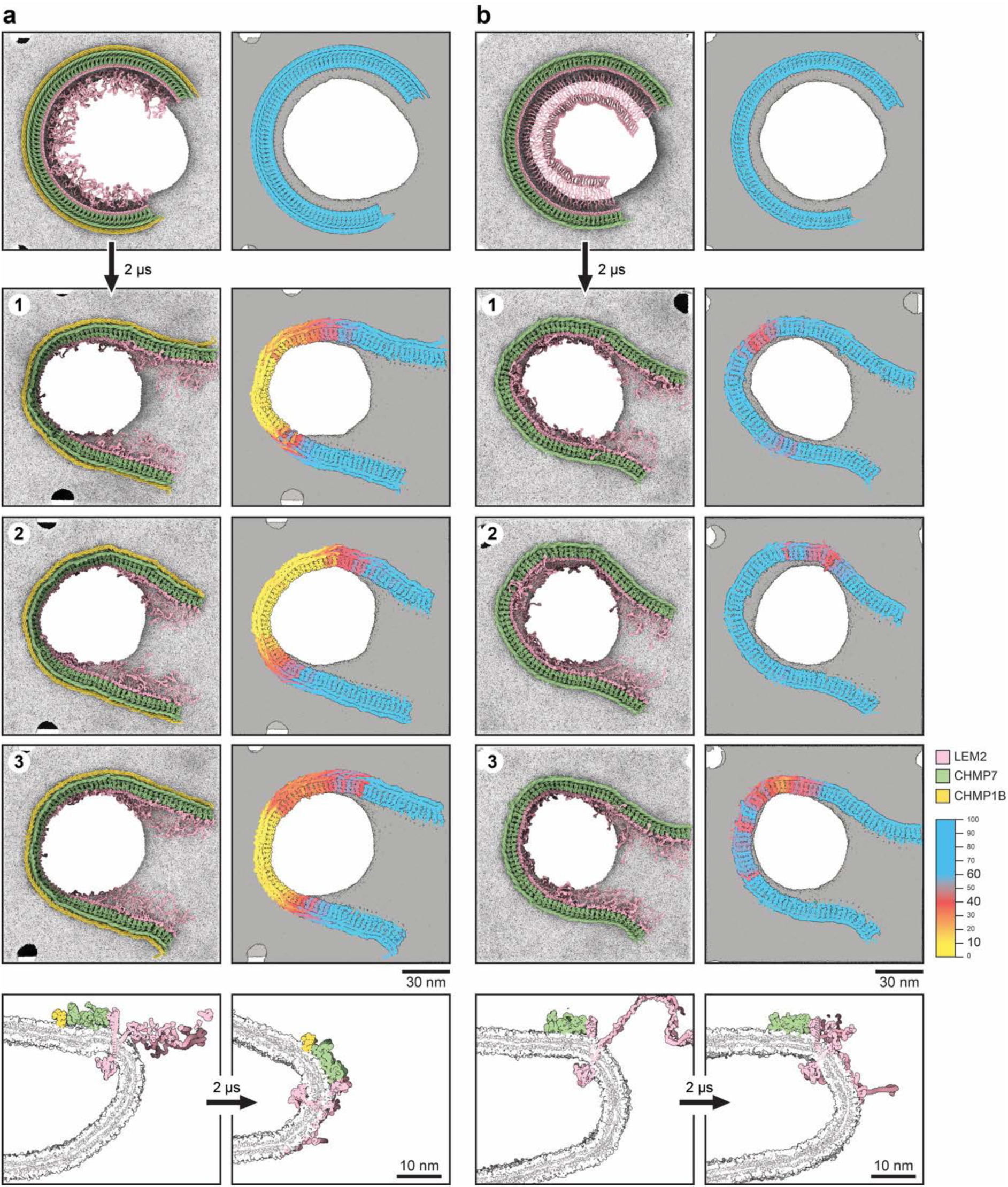
Tilting of LEM2-ESCRT-III filaments during membrane-pore simulations. Initially flat filaments of LEM2-CHMP7-CHMP1B **(a)** and LEM2-CHMP7 **(b)** were simulated on membrane pores using coarse-grained MD for 2 µs. For each system, three independent simulations were performed for each system (1–3). Colors on the right indicate the orientation of individual subunits relative to the membrane normal, quantified as the angle between the z-axis a vector connecting the centers of the CHMP7 WH1 and WH2 domains. In the presence of CHMP1B, the more rigid LEM2-CHMP7-CHMP1B filament accommodates the increasing pore curvature by tilting out of the membrane plane and moving toward the pore neck. In the absence of CHMP1B, the more flexible LEM2-CHMP7 filament adapts to the changing pore geometry primarily through in-plane deformation and remains predominantly flat at the outer region of the pore.

**Supp. Figure 11:**
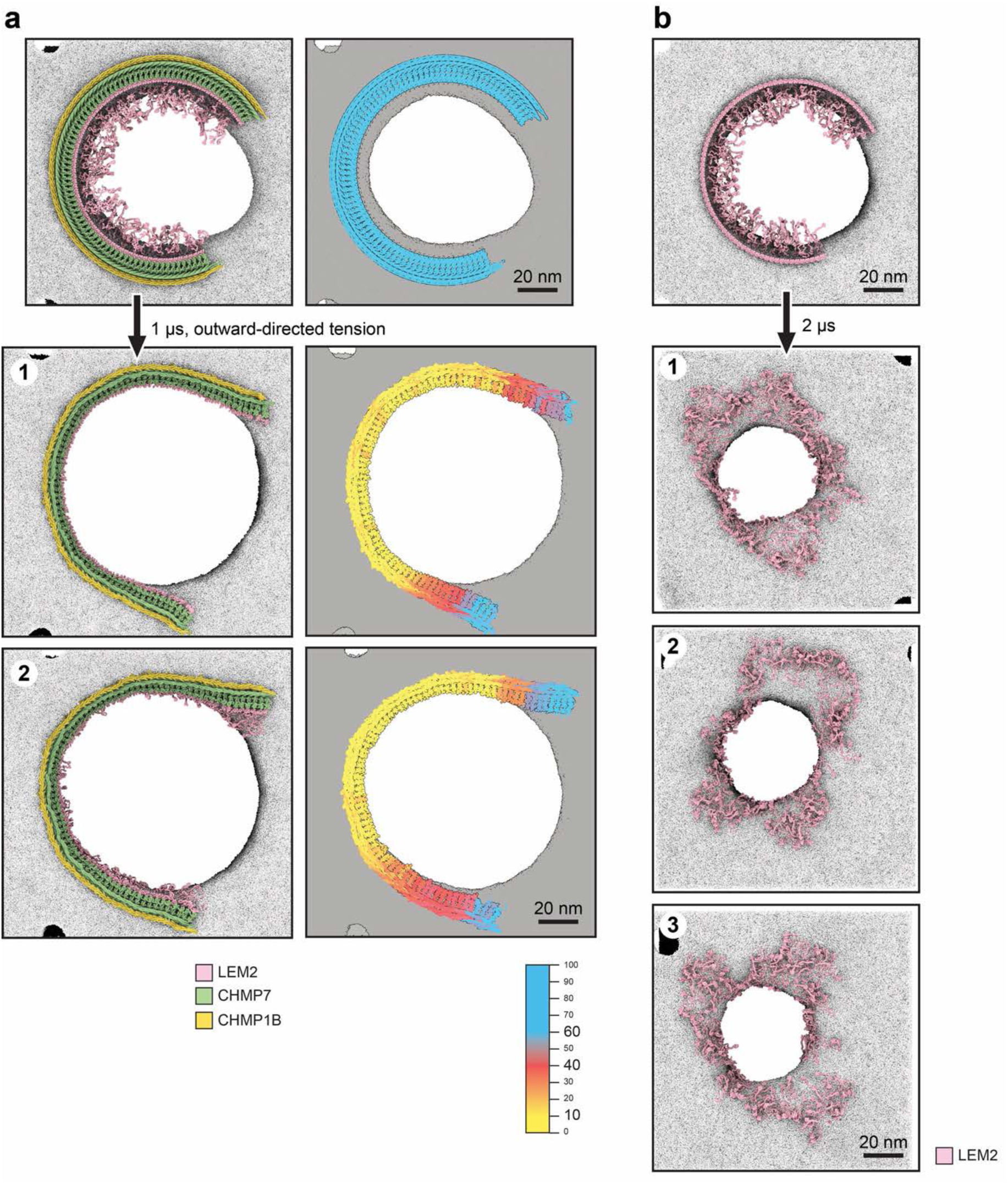
Applied lateral tension on LEM2-CHMP7-CHMP1B pore and LEM2-only pore simulations. **a)** Outward-directed tension was applied to a double-membrane pore containing a LEM2-CHMP7-CHMP1B filament for 1 µs. Two independent simulations are shown (1,2). Colors indicate the orientation of individual subunits relative to the membrane normal, quantified as the angle between the z-axis and a vector connecting the centers of the CHMP7 WH1 and WH2 domains. Under outward tension, the filament remains associated with the pore and adapts its orientation to the expanded membrane geometry. **b)** Coarse-grained MD simulations of a hypothetical highly curved LEM2-only strand positioned on a double-membrane pore for 2 µs. Three independent simulations are shown (1–3), illustrating the behavior of the LEM2 polymer in the absence of CHMP7 and CHMP1B.

**Supp. Figure 12:**
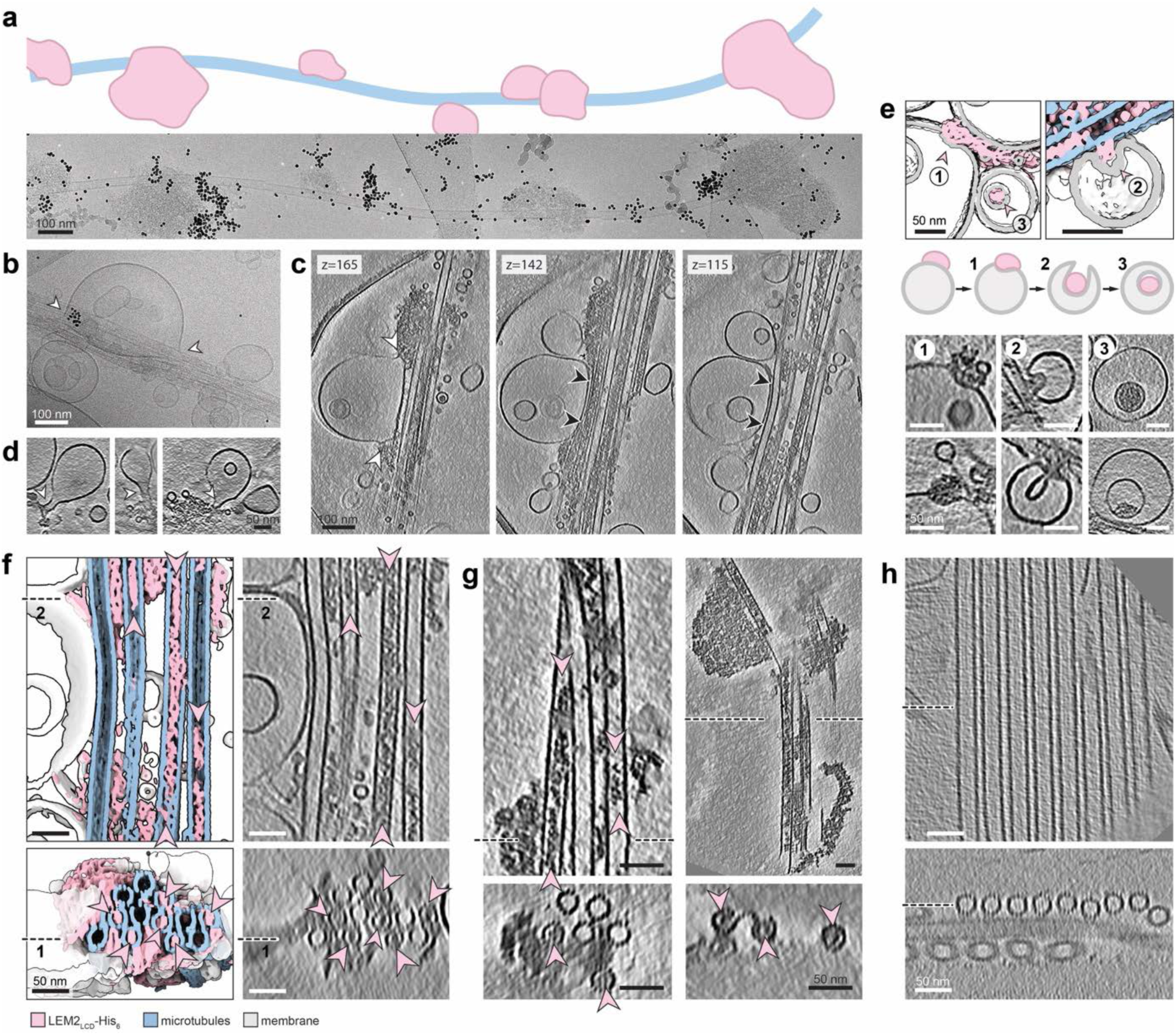
LEM2couples microtubules to membranes and accesses the microtubule lumen. **a)** Projection image of a microtubule, decorated with droplets of LEM2_LCD_. **b)** Projection image of a microtubule bundle and Ni-NTA vesicles, attached via LEM2_LCD_-His_6_. **c)** Tomographic slices of a microtubule bundle and Ni-NTA vesicles, attached via LEM2_LCD_-His_6_. The membrane is stretched (indicated by white arrow heads in **b** and **c**) and brought in tight contact to the microtubule (black arrow heads in **c**), leaving a separation of ∼3 nm. The microtubule is locally bent at sites of close membrane contact. **d)** Droplets of LEM2_LCD_-His_6_ condensates deform vesicles and generate narrow membrane necks. Small vesicles of ∼10–15 nm diameter are frequently observed in close proximity to these necks, consistent with membrane budding from these sites. **e)** LEM2_LCD_-His_6_ condensates interact with Ni-NTA vesicles and are associated with a series of membrane geometries ranging from shallow indentations (1) to deeper invaginations (2) and small condensate-filled intraluminal vesicles (3) **f,g)** LEM2_LCD_ (with His_6_-tag in **f**) enters the lumen of microtubules (**f**: in the presence of Ni-NTA vesicles); pink arrow heads indicate LEM2 density within the lumen. **h)** Control: microtubules without LEM2_LCD_ show a void lumen.

**Supp. Figure 13:**
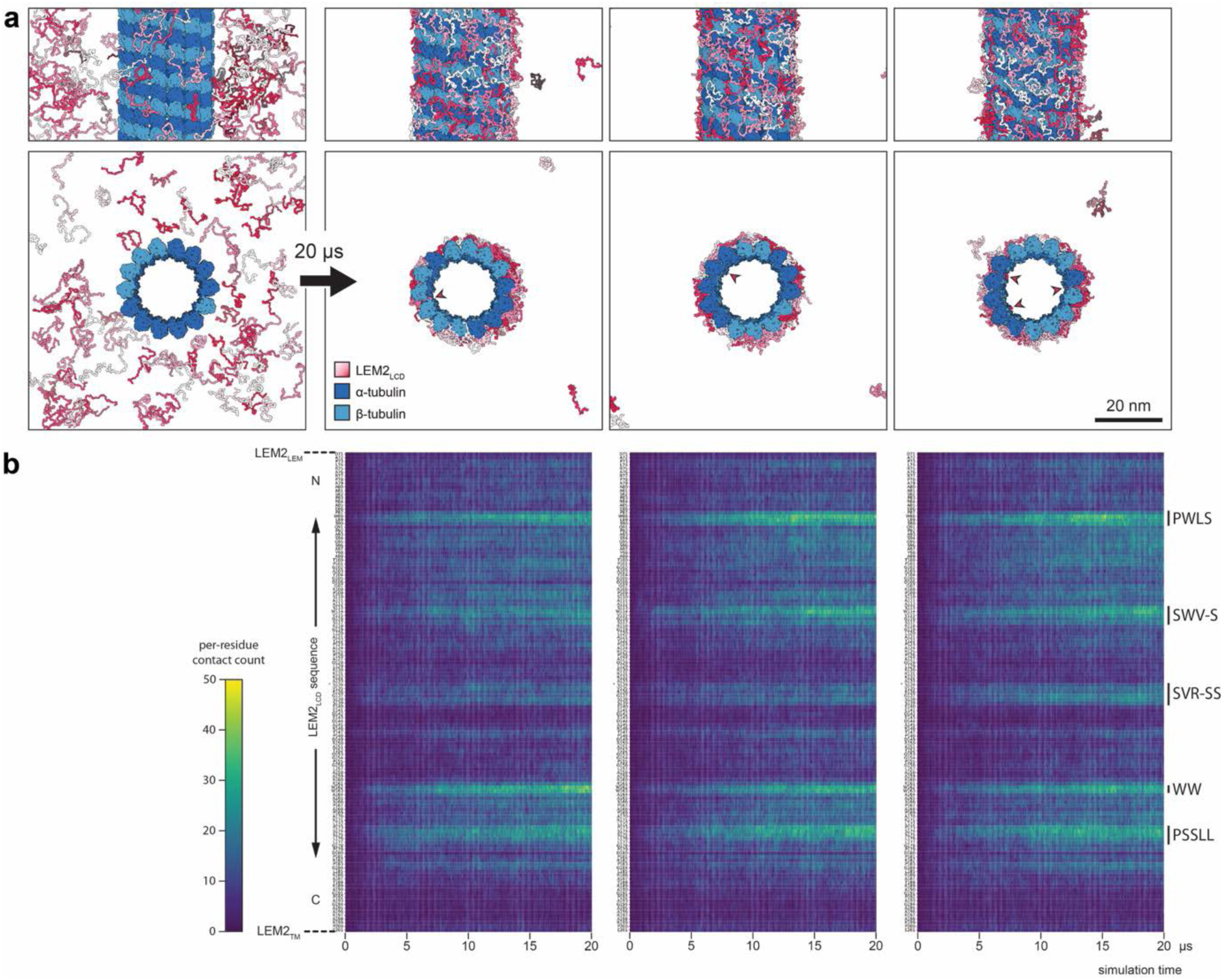
Interaction between LEM2_LCD-ΔLem_ and microtubules in coarse-grained MD simulations over 20 µs. **a)** Start (left) and end points of three independent simulations. 96 copies of LEM2_LCD-ΔLem_ were randomly placed in a simulation box of dimensions 80 x 80 x 42 nm and simulated together with a microtubule in the center. Five LEM2_LCD-ΔLem_ copies were observed to start entering the microtubule lumenal space (pink arrow heads). **b)** Interaction profiles of the LEM_LCD-ΔLem_ sequence, shown for each simulation replicate. Counted were residues that came into a distance radius of 6 Å with microtubules and averaged among the 96 copies of LEM2_LCD-ΔLem_. Residues of frequently interacting patches are highlighted on the right.

**Supp. Figure 14:**
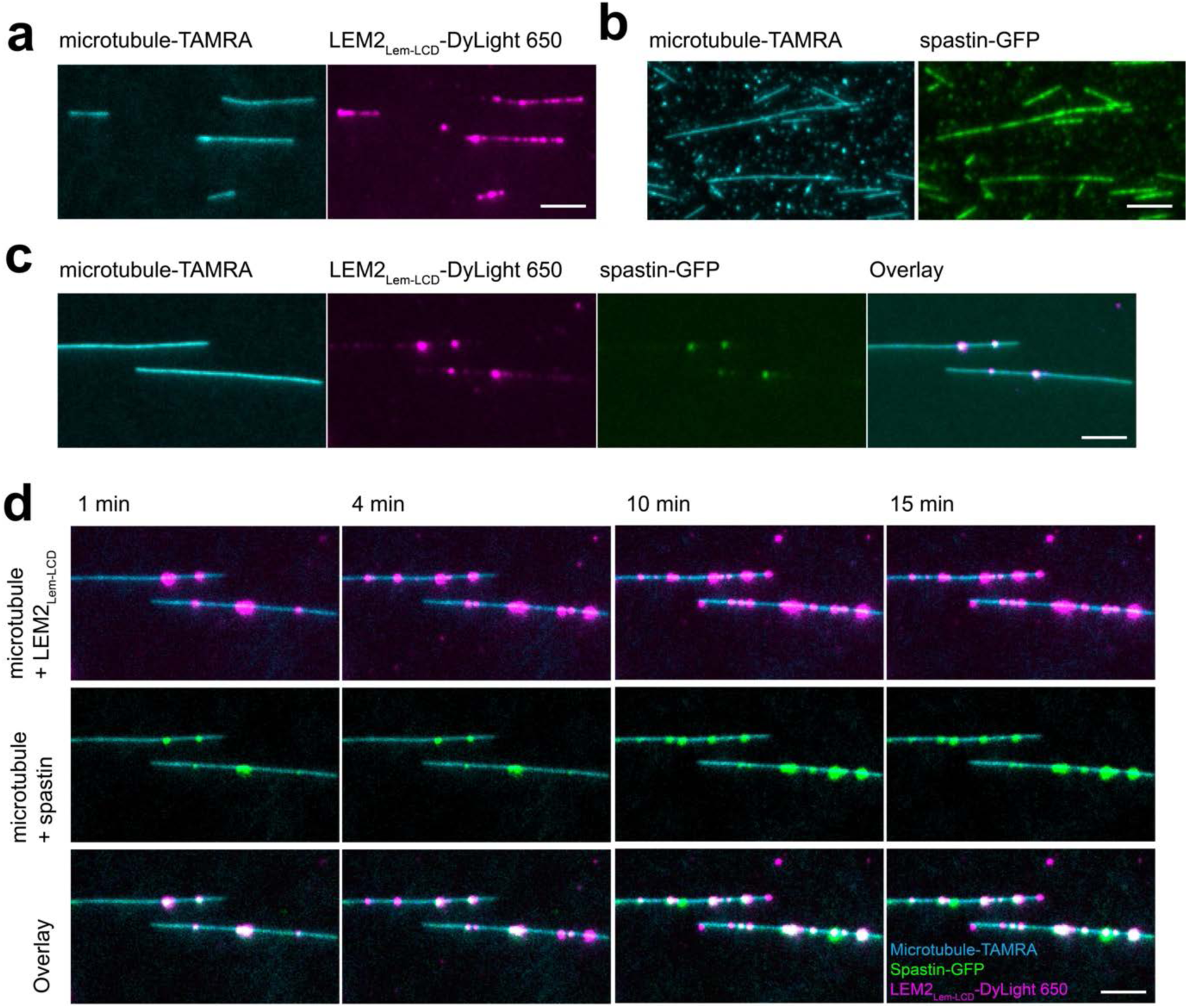
LEM2_LCD_ droplets and spastin colocalize on microtubules. Microtubules recruit **(a)** LEM2, **(b)** spastin. **c)** LEM2 and spastin colocalizes on microtubules. **d)** LEM2 droplets accumulate on microtubules with time, enriching spastin within the droplets. Scale bar, 5 μm.

**Supp. Figure 15:**
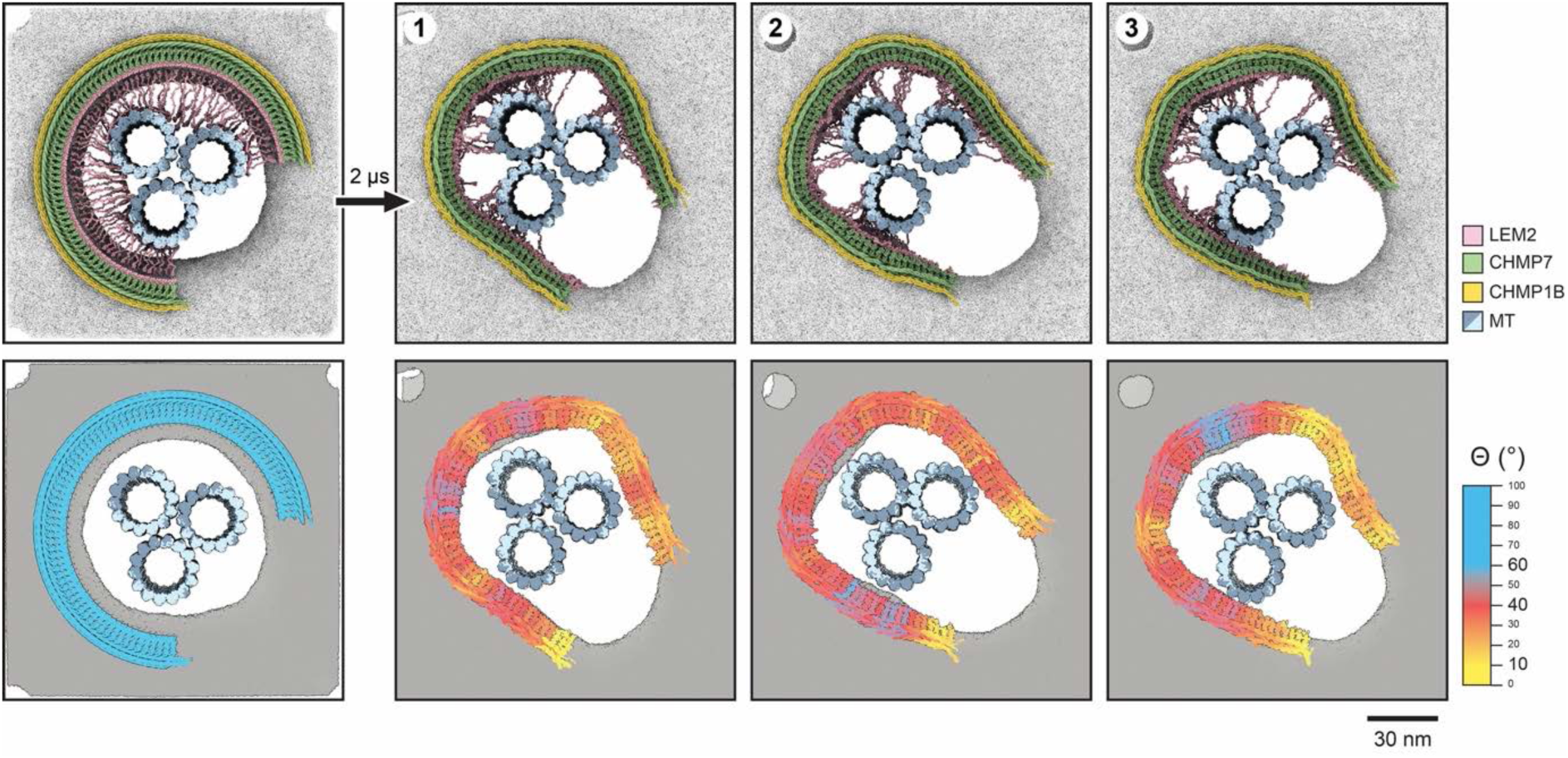
Tilting of an LEM2-ESCRT-III filament during LEM_LCD_-microtubule mediated interactions in membrane-pore simulations. An ¾ ring of a flat LEM2-CHMP7-CHMP1B filament was simulated on a membrane pore using coarse-grained MD for 2 µs. Three independent simulations were performed (1–3). Colors in the bottom panels indicate the orientation of individual subunits relative to the membrane normal, quantified as the angle Ɵ between the z-axis and a vector connecting the centers of the CHMP7 WH1 and WH2 domains. The filament tilts toward the pore neck while LEM2_-LCD_ forms extensive contacts with the microtubules. To facilitate initial interactions between LEM2_LCD_ and the microtubules and prevent premature adsorption of LEM2_LCD_ onto the membrane, LEM2_LCD_ was initialized in an extended configuration oriented toward the microtubules.

**Supp. Figure 16:**
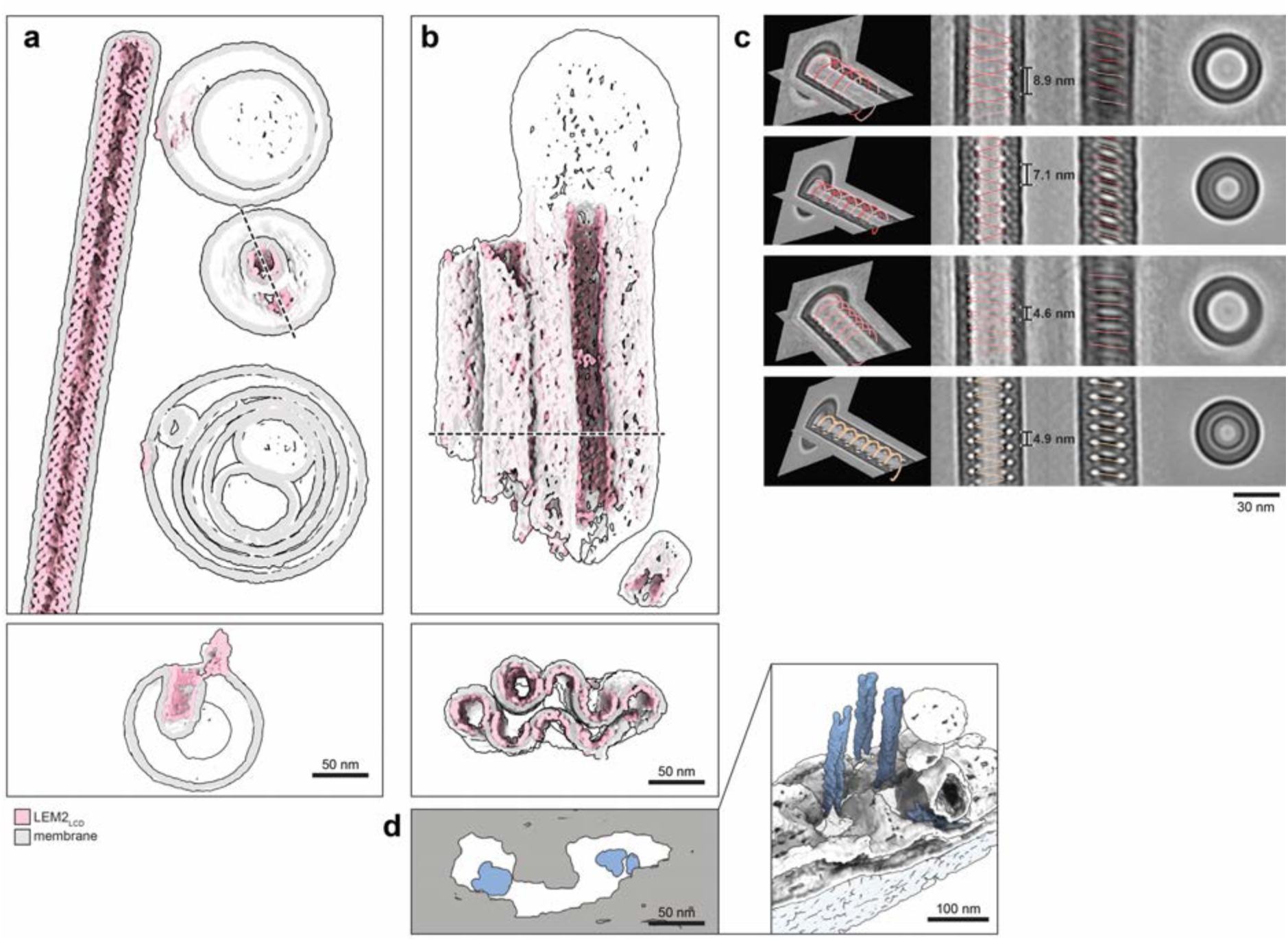
Negative-curvature membrane tube formation by LEM2_WH_ on vesicles (in the absence of CHMP7 and CHMP1B). **a)** Full tube and peripheral ¼-tubular filaments on vesicles. Indicated cross-section (bottom) showing a tubular invagination. **b)** Sheet-like structures, forming from ½, ¾ and full tubes. **c)** Subtomogram averages of LEM2_WH_ tubes. Tubes have a different helical symmetry than in the presence of CHMP7 and CHMP1B (see Supp. Fig. 17). **d)** Comparison of terminal pore geometry with membrane tubes generated by LEM2_WH_. Right: view of the terminal pore from the anaphase tomogram shown in Supp. Fig. 2. Left: Top view of the pore, matching the cross-view of negative membrane curvatures generated by LEM2_WH_ from panel **b**.

**Supp. Figure 17:**
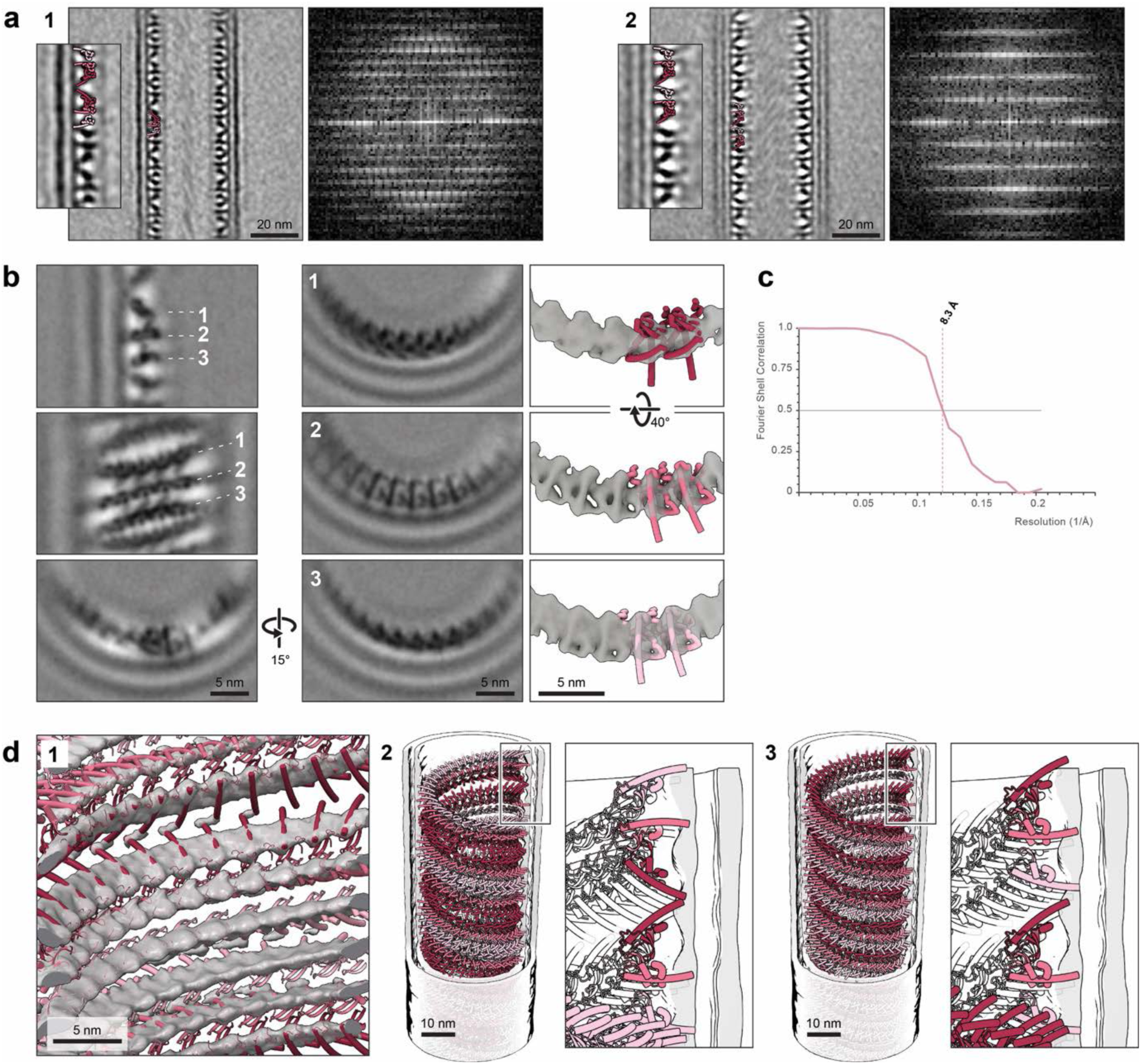
LEM2_WH_ forms negatively curved membrane tubes with regular diamter with two predominant helical architectures. Samples contained LEM2_WH_, CHMP7 and CHMP1B; negative membrane tubes consist only of LEM2_WH_. **a)** Subtomogram averages of individual LEM2_WH_ tubes with antiparallel (1) and parallel (2) two-start helical architectures. Each of the two helical starts comprises three LEM2_WH_ strands with distinct orientations relative to the membrane. Right panels show layer-line patterns in reciprocal space, which aided assignment of the helical architecture in noisy averages of individual tubes. In the antiparallel architecture, the axial repeat is effectively doubled relative to the parallel architecture, producing additional layer lines at intermediate positions. **b)** Subtomogram average of antiparallel LEM2_WH_ tubes shown in central sections (left), sections peripheral to the helical axis (center), and isosurface representations with pairs of rigid-body-fitted AlphaFold-derived LEM2_WH_ models (right) **c)** Fourier shell correlation (FSC) curve indicating a resolution of 8.2 Å at an FSC cutoff of 0.5. **d)** Helical architecture of LEM2WH tubes shown as atomic models fitted into the density (1) and as cut-open representations of the antiparallel (2) and parallel (3) assemblies.

## Methods

### Purification of proteins

#### LEM2_WH_

Human LEM2_WH_ (residues 395–503) was expressed in BL21-(DE3)-RIPL cells and purified as previously described as His_6_-Sumo fusion protein (8). In short, collected cells were resuspended in lysis buffer (40 mM HEPES, pH 8.0, 350 mM KCl, 5% glycerol, 10 mM imidazole, 2 μg/ml DNase I, 5 mM β-mercaptoethanol, lysozyme and protease inhibitors) at 5 ml per gram of cell pellet. Cells were lysed by sonication on ice, and the lysate was clarified by centrifugation (10,000g, 30 min, 4 °C). The clarified lysate was incubated with Ni-NTA agarose resin (Qiagen; 4 ml bed volume) for 1.5 h at 4 °C. The resin was washed with 500 ml wash buffer (40 mM HEPES, pH 8.0, 350 mM KCl and 5% glycerol), and bound protein was eluted with five column volumes of wash buffer supplemented with 350 mM imidazole. The eluate was spin-concentrated (Vivaspin 20, 3 kDa MWCO, PES) to approximately 5 ml and dialysed overnight at 4 °C against wash buffer with 1 mM DTT. His_6_-Sumo-LEM2_WH_ was further purified by size-exclusion chromatography using a Superdex 75 16/60 column (GE Life Sciences) equilibrated in the same buffer. His_6_-Sumo-LEM2_WH_-containing fractions were pooled, spin-concentrated (Vivaspin 6, 3 kDa MWCO, PES), aliquoted, snap-frozen in liquid nitrogen and stored at −80 °C. The His_6_-Sumo tag was cleaved for 30 min at room temperature immediately before use using His_6_-Ulp1 protease. Before use, thawed protein was cleared of aggregates by centrifugation.

#### LEM2_Lem-LCD_

Human LEM2_Lem-LCD_ (residues 1–208) was expressed as a SUMO-LEM2_Lem-LCD_-HRV3C-His_6_ fusion and purified as previously described for purification of LEM2_NTD_ (44). All purification steps were performed at 4 °C. The protein was expressed in BL21 T7 Express pRare cells. Harvested cells were resuspended in lysis buffer (30 mM HEPES, pH 7.4, 800 mM KCl, 5% glycerol, 20 mM imidazole, 1 mM MgCl_2_, Benzonase and protease inhibitors) at 5 ml per gram of cell pellet and lysed by high-pressure homogenization. The lysate was clarified by centrifugation (10,000 g, 30 min, 4 °C) and applied to Ni-NTA resin. The resin was washed sequentially with wash buffer (30 mM HEPES, pH 7.4, 800 mM KCl, 5% glycerol) containing 20, 30 and 50 mM imidazole, and bound protein was eluted with wash buffer supplemented with 300 mM imidazole. Protein-containing fractions were pooled and concentrated, and the Sumo tag was cleaved with His_6_-Ulp1 protease during overnight dialysis against 30 mM HEPES, pH 7.4, 400 mM KCl and 5% glycerol. Cleaved LEM2_Lem-LCD_-His_6_ was further purified by size-exclusion chromatography using a HiLoad Superdex 200 pg 16/600 column equilibrated in the same buffer. LEM2_Lem-LCD_-His_6_-containing fractions were pooled, concentrated, aliquoted, snap-frozen in liquid nitrogen and stored at −80 °C. Before use, thawed protein was cleared of aggregates by centrifugation.

#### CHMP7

Full-length human CHMP7 was expressed in BL21-(DE3)-RIPL cells as a His_6_-SUMO fusion and purified as previously described (8). Bacterial cell pellets (20–30 g) were resuspended in lysis buffer (50 mM Tris, pH 8.0, 800 mM KCl, 20 mM imidazole, 10% glycerol, 5 mM β-mercaptoethanol, 2 μg/ml DNase I and protease inhibitors, Lysozyme) at 5 ml per gram of cell pellet. Cells were lysed by sonication on ice, and the lysate was clarified by centrifugation (10,000 g, 30 min, 4 °C). The clarified lysate was incubated with Ni-NTA agarose resin (Qiagen; 4 ml bed volume) for 1.5 h at 4 °C. The resin was washed with 500 ml wash buffer (50 mM Tris, pH 8.0, 800 mM KCl, 10% glycerol) supplemented with 20 mM imidazole and eluted with five column volumes of wash buffer supplemented with 350 mM imidazole. Eluted protein was dialyzed on at 4C against gel-filtration buffer (50 mM Tris, pH 8.0, 800 mM KCl, 1 mM DTT and 5% glycerol). His_6_-Ulp1 protease was added during dialysis to cleave the His_6_-Sumo tag, and dialysis was continued overnight. The cleaved sample was incubated with Ni-NTA agarose resin to remove His_6_-Sumo, His_6_-Ulp1 and Ni-NTA-binding contaminants. The flow-through was concentrated to approximately 5 ml (Vivaspin 20, 30 kDa MWCO, PES) and monomeric CHMP7 was isolated by size-exclusion chromatography using a Superdex 75 16/60 column (GE Life Sciences) equilibrated in gel-filtration buffer. CHMP7-containing fractions were concentrated, aliquoted, snap-frozen in liquid nitrogen and stored at −80 °C. Before use, thawed protein was cleared of aggregates by centrifugation.

#### CHMP1B

Human CHMP1B was purified as previously described (27) using an N-terminal His_6_-SUMO fusion. His_6_-SUMO-CHMP1B was expressed in LOBSTR-BL21(DE3) cells in ZYP-5052 auto-induction medium (45). Cells were harvested, frozen at −80 °C and subsequently resuspended in lysis buffer (50 mM Tris, pH 8.0, 500 mM NaCl, 10 mM imidazole, 1 mM DTT and 5% glycerol). All subsequent purification steps were performed at 4 °C. Lysozyme was added and cells were lysed by sonication. The lysate was clarified by centrifugation (30,000 g, 1 h), filtered through a 0.45-μm membrane and incubated with Ni-NTA resin (Qiagen) for 1 h. The resin was washed with 500 ml wash buffer (50 mM Tris, pH 8.0, 500 mM NaCl, 1 mM DTT), and bound His_6_-Sumo-CHMP1B was eluted with wash buffer supplemented with 400 mM imidazole. His_6_-Ulp1 protease was added to the eluate, which was dialysed overnight at 4 °C against cleavage buffer (20 mM Tris, pH 8.0, 150 mM NaCl and 1 mM DTT). The cleaved sample was passed over Ni-NTA resin a second time to remove His_6_-Sumo, His6-Ulp1 and residual uncleaved fusion protein. CHMP1B was further purified by size-exclusion chromatography using a Superdex 75 16/60 column (GE Healthcare Life Sciences) equilibrated in 20 mM Tris, pH 7.4, 150 mM NaCl and 1 mM DTT. Purified proteins were aliquoted, snap-frozen in liquid nitrogen and stored at −80 °C. Before use, thawed protein was cleared of aggregates by centrifugation.

### LEM2_WH_-ESCRT-III *in vitro* reconstitutions

#### Generation of small unilamellar vesicles (SUVs)

SUVs were generated from an NE-like membrane mixture as suggested previously (46). 54.5% POPC, 15% cholesterol, 13.4% Liver PI, 8% POPE, 5.38% POPA, 3.57% DOPS and 1.5% Rhod PE were mixed to a final total lipid concentration of 1 mM. Corresponding mole percent of lipids were transferred to a glass vial containing 200 µL chloroform and dried in the vial using a nitrogen stream and desiccator, and subsequently resuspended in 500 µL 20 mM HEPES, 75 mM NaCl, 1 mM DTT, pH 8 to generate a final concentration of 1 mM. The vial was incubated for 1 h at 37°C on a shaker and subjected to 10 freeze-thaw cycles between 1 min in liquid nitrogen and 5 min in 42 °C water. Lipid aggregates from the resulting mixture were removed by centrifugation for 30 min, 17,000 g at room temperature and the supernatant was collected. SUVs were generated by 21 cycles of extrusion through a membrane with 200 µL pore diameter in an Avestin LiposoFast liposome extruder, aliquoted and stored at −80°C.

#### Preparation of detergent-solubilized lipids

The stock solution of detergent-solubilized lipids used in this study contained 2 mg/mL DOPS in 40 mM CHAPS, 40 mM HEPES, 50 mM NaCl, pH 8.

#### LEM2_WH_-CHMP7-CHMP1B and LEM2_WH_-CHMP7 filament assembly with solubilized lipids

14 µM His_6_-Sumo-LEM2_WH_ and 13 µM CHMP7, with or without 13 µM CHMP1B, were mixed with 0.32 mg/mL DOPS/CHAPS in 20 mM HEPES, 500 mM NaCl, pH 8 and dialyzed (10,000 MWCO, Thermo Scientific) against 20 mM HEPES, 75 mM NaCl, 1 mM DTT, pH 8 at room temperature overnight. His_6_-Ulp1 was added at a ratio of 1:12.5 for 90 min. The filaments were centrifuged with 5,000 rpm for 10 min, the supernatant was removed, the pellet was washed once and resuspended in 20 mM HEPES, 75 mM NaCl, 1 mM DTT, pH 8 in equivalents of the original volume. Grids of these assemblies were prepared from a dilution of 1:10.

#### LEM2_WH_-CHMP7-CHMP1B assembly on membranes

6 µM His_6_-Sumo-LEM2_WH_, 4.6 µM CHMP7 and 4.6 µM CHMP1B were mixed with 0.5 mg/mL SUVs and Ulp1 at a ratio of 1:25 in 20 mM HEPES, 500 mM NaCl, pH 8 and dialyzed (10,000 MWCO, Thermo Scientific) against 20 mM HEPES, 75 mM NaCl, 1 mM DTT, pH 8 at room temperature overnight. Grids of these assemblies were prepared from a dilution of 1:7.

#### CHMP7 assembly on membranes

4.6 µM CHMP7 was mixed with 0.5 mg/mL SUVs in 20 mM HEPES, 500 mM NaCl, pH 8 and dialyzed (10,000 MWCO, Thermo Scientific) against 20 mM HEPES, 75 mM NaCl, 1 mM DTT, pH 8 at room temperature overnight. Grids of these assemblies were prepared from a dilution of 1:3.

#### LEM2_WH_ assembly on membranes

8 µM His_6_-Sumo-LEM2_WH_ was mixed with 0.5 mg/mL SUVs and Ulp1 at a ratio of 1:25 in 20 mM HEPES, 500 mM NaCl, pH 8 and dialyzed (10,000 MWCO, Thermo Scientific) against 20 mM HEPES, 75 mM NaCl, 1 mM DTT, pH 8 at room temperature overnight. Grids of these assemblies were prepared from the undiluted sample.

### LEM2_Lem-LCD_-microtubule *in vitro* reconstitutions

#### Generation of double-stabilized MTs

Microtubule assembly followed protocols described in (47,48) and all microtubule-containing assays were performed at room temperature. 40 µL of microtubule seed mix was prepared from 4.5 µM Tubulin with 1 mM GMP-CPP, 1 mM MgCl_2_, in BRB80 buffer (80 mM PIPES, 1 mM MgCl_2_, 1 mM EGTA, pH 6.9) and incubated on ice for 5 min, followed by an incubation of 30 min at 37°C. The seed mix was centrifuged at 13300 rpm for 15 min at 25 °C. The pellet of short microtubules was resuspended in 39.5 µL BRB80, 1 mM GMP-CPP and 1 mM MgCl_2_. Additional 0.5 µL of 40 µM tubulin was added and incubated for 24 h at 37 °C. This elongation mix was centrifuged again and the pellet resuspended in 20 µL BRB80, supplemented with 10 µM taxol and stored at room temperature.

#### LEM2_Lem-LCD_-MT assemblies

8 µM LEM2_Lem-LCD_ were mixed with 10% (v/v) microtubules in taxol-supplemented BRB80 and incubated in a tube for ∼30 min before freezing.

#### LEM2_Lem-LCD_-MT-SUV assemblies

8 µM LEM2_Lem-LCD_ were mixed with 10% (v/v) microtubules in taxol-supplemented BRB80 and incubated for 10 min. This mixture was applied to a cryo-TEM grid, incubated for 2 min, mixed with 4% Ni-NTA SUVs in a ratio of 1:1 and incubated for another 5 min before blotting and freezing.

### Preparation of cryo-TEM grids by plunge freezing

TEM gold grids with holey carbon support film (Quantifoil Micro Tools GmbH, Cu 300 mesh R1.2/1.3) were cleaned with chloroform for ca. 30 min and glow-discharged from both sides (PELCO, easyGlow). For plunge freezing, an EM Grid Plunger (Leica) was used. At 18°C, 3 µL diluted samples of *in situ* reconstitutions were applied to the grid and mixed with 1 µL sonicated 10-nm colloidal gold particles (BBI solutions). Blotting was performed from the back for 1 sec, the grid was plunged into liquid ethane (Air Liquide) and stored in liquid nitrogen until data acquisition.

### Data acquisition

Screening of samples and tilt series acquisition (predominantly for analysis by segmentation) were performed on a Titan Halo (Thermo Scientific) cryo-transmission electron microscope operating at 300 kV and equipped with a Quantum-K2 imaging filter (Gatan). For any samples containing LEM2_WH_, CHMP7 and/or CHMP1B, a magnification of 105,000x (1.537 Å/px) was used; for samples containing LEM2_Lem-LCD_ and microtubules, a magnification of 61,000x (2.303 Å/px) was used. Tomographic tilt series were acquired using a bidirectional tilt scheme, starting at 18 or 20°, with maximal tilt ranges between ±44 and ±60° and tilt increments of 2, 3 or 4°. Digital Micrograph software (Gatan) was used to tune the imaging filter, and SerialEM software versions 4.0.3 to 4.2.0 (49) were employed for the automated acquisition of tilt series in low-dose mode.

Tilt series for high-resolution reconstructions of the samples LEM2_WH_-CHMP7-CHMP1B in solubilized lipids and LEM2_WH_-CHMP7-CHMP1B on small unilamellar vesicles were acquired at a Titan Krios G4 (Thermo Scientific) cryo-transmission electron microscope operating at 300 kV equipped with a cold FEG, a Selectris X imaging filter using a slit width of 10 eV, and a Falcon 4 direct electron detector recording each tilt image as a movie with 10 frames at a magnification of 105,000x, corresponding to a pixel size of 1.223 Å. A dose-symmetric tilt scheme was used with a maximal tilt range of ±45° and an increment of 3°. Each tilt series had a total dose of 90 e^−^/Å^2^. For LEM2_WH_-CHMP7-CHMP1B in solubilized lipids, the targeted defocus was set to between −1 and −2 µm, and for LEM2_WH_-CHMP7-CHMP1B on small unilamellar vesicles, a range of −1 to −3 µm was used. SerialEM software versions 4.0.6 and 4.0.14 (49) were used for automated acquisition of tomographic tilt series in low-dose mode.

A summary of acquired cryo-electron tomograms is provided in Table S1.

**Table S1:**
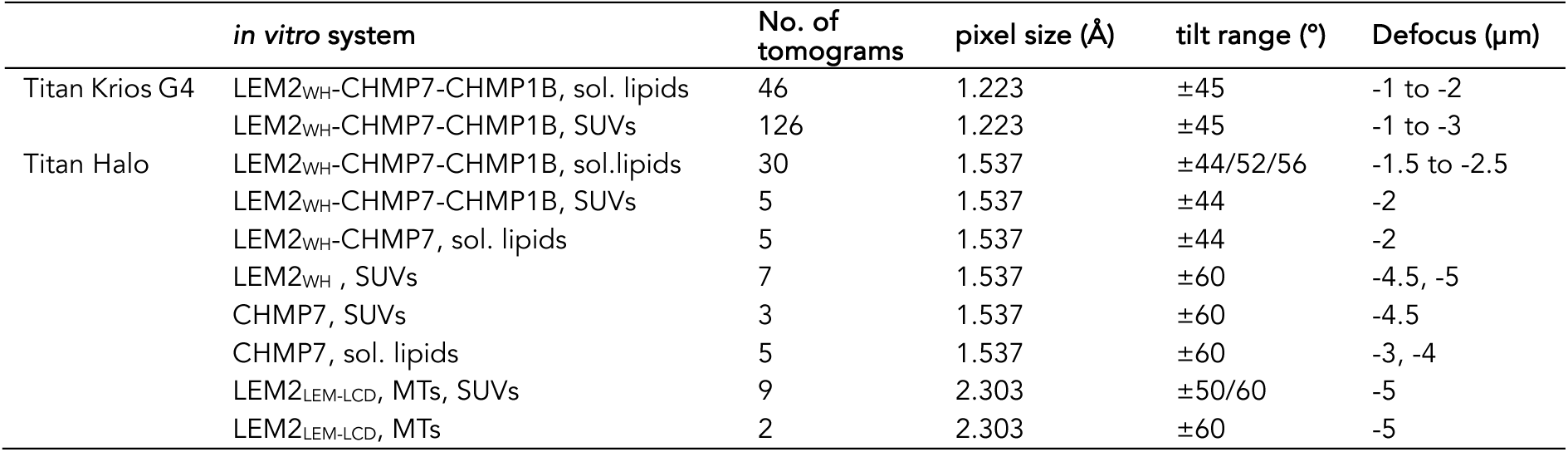
Acquired cryo-electron tomograms for different *in vitro* systems.

### Tomographic reconstruction

Motion correction of tilt series frames was performed using MotionCorr (50). Tomograms were reconstructed with ETOMO from IMOD version 4.12.50 (51) using fiducial markers for alignment, dose-weight filtering, 3D-CTF correction by phase flipping and weighted back-projection. For visualization and segmentation, tomograms that where acquired with a tilt range of ±50° or higher were denoised and more isotropically reconstructed using IsoNet (52).

### Segmentation

Segmentation of tomograms was performed with DragonFly version 2022.2.0.12227 (Object Research Systems, 2022; www.theobjects.com/dragonfly). Room-temperature tomograms of anaphase cells had pixel sizes of 7.67 and 4.51 Å and were binned 2x before segmentation. For most cryo-electron tomograms, IsoNet-treated bin8 tomograms were used, except for samples of LEM2_WH_-CHMP7-CHMP1B + SUVs, for which the limited tilt range of ±45° precluded this treatment and NAD-filtered tomograms were used instead. Initial segmentations were manually done using local-Otsu based thresholding on a small training area of around 400×400×50 px, on which a U-Net was trained. The segmentation was then predicted on the full tomogram and manually cleaned and refined. Segmentations were exported as 2D-tiff files, converted into mrc files with IMOD and imported into ChimeraX (53) for visualization.

### Curvature analysis

The curvatures of LEM2_WH_-CHMP7-CHMP1B and LEM2_WH_-CHMP7 filaments with soluble lipids were measured from the manually modeled contours that were also used as starting models for subtomogram averaging. Their local radii were computed in 3D on bin8 tomograms with the program “imodcurvature”, using a window length of 100 px.

### Subtomogram averaging

#### LEM2_WH_-CHMP7-CHMP1B filaments with solubilized lipids

Particle picking and initial alignment was performed using PEET version 1.18.0 (54). On bin8 tomograms (pixel size 9.784 Å), open contours were manually modeled following filaments on their bicellar lipid density within their C2 symmetry center. Particle points were then placed every 3 nm, resulting in a total number of 181,369 initial particles. Using a loose mask, particles were pre-aligned allowing for ±180° rotation around the filament axis, while limiting the other axes to ±21° and the shift along the filament to 1 px. The resulting alignment was then transferred to RELION-4.0 (55) and repeated on bin8 including 9 repeats within the mask, before moving to bin4 data and shrinking the mask to 7 repeats. Since the filaments appeared flexible in their curvature in tomograms and the antiparallel strands showed heterogenous appearance in cross views by eye, we did not apply C2 symmetry to the whole structure. Instead, the particle center was shifted into one strand, specifically the WH2 domain of CHMP7, and all particles were rotated by 180° to align subunits from both antiparallel strands, resulting in a total of 362,740 particles. Subsequently, the binning was reduced to 2 and alignment was iterated on a segment containing only 5 repeats. Particle coordinates were loaded into ArtiaX (56) and particles poorly following their filament were erased. Duplicate particles were removed. Classification with four classes was performed, keeping only particles of the highest-quality class, resulting in a final total of 38,278 particles. CTF correction and frame alignment were refined, before the final alignment was performed. The resolution was determined on a segment including 5 repeats of each LEM2_WH_ and CHMP7. See table S2 for refinement statistics.

#### Helical LEM2_WH_ membrane tubes (derived from the sample containing LEM2_WH_, CHMP7, CHMP1B and SUVs)

Particle picking and alignment was performed using PEET (54). On bin8 tomograms (pixel size 9.784 Å), open contours were manually modeled following the center of a total of 188 negatively curved membrane tubes. Particle points were then placed every 10 nm within the tube center, roughly following the height of the helical pitch, resulting in 3255 points. For each tube, an individual average was generated, using a central single particle as reference and a cylindrical loose mask including the complete tube structure over a length of 50 nm. These intra-tube averages where then manually classified in antiparallel or parallel tubes using their layer lines within their power spectra and overall visual appearance. This resulted in 73 antiparallel and 29 parallel tubes; the remaining 86 tubes were excluded due to ambiguity or different appearance. Antiparallel tubes were processed further. Their points were copied 36 times and incrementally rotated by 10° around their tube axis, resulting in 51588 particles. Alignment was performed allowing for shifts of 10 nm along their tube axis, while limiting rotations to 2°. This allowed particles to register within the helical rise. Next, particles were loaded in ArtiaX (56). Some tubes revealed segments that showed parallel instead of antiparallel alignment and those particles, as well as particles poorly following their helical structure, were erased, resulting in 17707 particles from 36 tubes. To account for their C2 symmetry without having to include the large volume of 6 individual LEM2_WH_ strands, particles were duplicated and rotated by 180° to align subunits from both antiparallel triple-strands. Then, the particle center was shifted from the tube center onto the center of one LEM2_WH_ triple-strand, and the alignment was brought to bin4.

The average at this point did not show repetitive subunit features laterally along the helical strands. In our hands, alignments with RELION did not converge on a regular repeat but consistently resulted in pronounced overfitting of noise. We therefore continued the alignment in PEET. To enhance subunit resolution, we returned to our most regular tube and manually selected a single particle at the topmost point of the flat-lying tube, where the subunit and its neighboring subunits were well resolved as their resolution was minimally affected by the missing wedge. An intra-tube average was generated using this particle as a reference, which showed exceptional resolution of subunit features. This intra-tube average was used as reference to align the remaining bin-4 particles within a mask spanning across 10 lateral subunits of a triple-strand. The data were brought to bin 2 for final alignment and resolution estimation. See table S2 for refinement statistics.

### Model building

#### LEM2_WH_-CHMP7-CHMP1B filaments with solubilized lipids

The final subtomogram average density contained 3 well-resolved repeats of single-stranded LEM2_WH_-CHMP7-CHMP1B. However, the elongated open ESCRT-III domains of CHMP7 and CHMP1B extended beyond the borders of the reconstructed volume, and to generate a density following their protein chains, 4 density copies were placed along the strand, overlapping by one repeat and fitted with ChimeraX fit-in-map function. Densities of the neighboring subunits were masked away to generate individual subunit densities for each protein. Each 12 copies of the three subunit densities were placed along the previously modeled filament strand using a helical symmetry with an angle of 3.365° and a rise of 9.44 Å. All 36 density maps were subsequently merged with the ChimeraX vop max command and used as a template map for atomic model refinement later.

The initial structure for LEM2_WH_ was obtained from AlphaFold (57,58) (AF-Q8NC56-F1) and we removed the Lem, LCD, lumenal and transmembrane domains to include only AA 395-503, corresponding to the biochemical construct used in our assembly assays. The predicted structure showed a good fit to our density.

CHMP7_WH1-WH2_ (AA 1-234) was taken from AlphaFold (AF-Q8WUX9-F1) as well, and the entity of both domains (except the NTD AA 1-19 and the membrane binding loop AA 113-144, which were not resolved) showed a good fit to our density, too. Its ESCRT-III domain, instead, was predicted by AlphaFold to have a closed conformation. Instead, we generated a homology-model (59) performed against the open ESCRT-III structure of the Snf7 core domain (PDB 5FD7) (22) and fused this to the WH1-WH2 domains. We removed N-terminally AA 1-19 and C-terminally ESCRT-III helix 6 (AA 394-453), which were not resolved in the density.

The initial structure for CHMP1B was taken from PDB 6TZ9 (27). Our density only showed AA 1-130 (until the first half of helix 4) and we removed AA 131-199.

Next, ESCRT-III helices of CHMP7 and CHMP1B were rotated to better follow the density by minimal angular adjustments within the elbow and loop regions between helices. Further refinement was performed with the ChimeraX plugin ISOLDE (60). Secondary structure elements were restricted in all models. The structures were first relaxed without map weight, before each protein chain was relaxed within its subunit density map and a map weight of 0.3. Structures were then placed into the 12-mer density map, helically symmetrized (3.365°, 9.44 Å), and dynamically refined with a map weight of 0.1. Clashes and rotamers were addressed systematically. Then, the central subunits of each protein were symmetrized again and refined over two more iterations before final symmetrization. See table S2 for model validation.

#### Adįusting the LEM2_WH_-CHMP7-CHMP1B filament for MD simulations on membrane pores

For MD simulations, LEM2_WH_ was extended to its full-length structure as obtained from AlphaFold (Supp. Fig. 4d). For CHMP7, we additionally included 1-19, but not 366-453. For CHMP1B, 131-199 remained truncated. The native cryo-ET structure contained a large pitch, which precluded filament placement on the flat surface of a double-membrane pore. To remove the pitch while maintaining lateral interactions, the filament was symmetrized planarly around an axis perpendicular to a plane peripheral to the helical rise (Supp. Fig. 9). The new lateral interactions and LEM2_FL_ were iteratively relaxed and symmetrized in ISOLDE.

For MD simulations including microtubules within the pore center, we ensured efficient initial contacts between LEM2_Lem-LCD_ and the microtubules. Therefore, we used the analysis from our previous simulations (Fig. 4b) to position the microtubule-interacting residues of LEM2_LCD_ facing the microtubules. Again, LEM2_Lem-LCD_ domains were relaxed using ISOLDE. In these simulations, three microtubules were included to fill out the available space in the pore center and preserve the filament structure’s diameter.

#### Helical LEM2_WH_ filaments in negatively curved membrane tubes

The subtomogram average density revealed repetitive subunits features for three LEM2_WH_ strands, however, the membrane-contacting parts of the elongated helix remained unresolved. Combined analysis of isosurfaces and slices allowed rigid fitting of the LEM2_WH_ domain, resulting in two parallel and one antiparallel strand. Small globular densities within the membrane layer helped to position the membrane-contacting helix (Supp. Fig. 17d). Between two antiparallel strands, the beta-sheet loops showed inter-strand interaction and further aided subunit placements. We measured a helical rise of −6.5 Å and a twist angle of 11.4°.

**Table S2:** Cryo-EM data collection, refinement and model validation.

|  | LEM2<sub>WH</sub>-CHMP7-CHMP1B | LEM2<sub>WH</sub> |
| --- | --- | --- |
| **Data collection and processing** |  |  |
| Magnification | 105,000 | 105,000 |
| Voltage (kV) | 300 | 300 |
| Electron exposure (e-/Å<sup>2</sup>) | 90 | 90 |
| Defocus range (µm) | -1 to -2 | -1 to -3 |
| Pixel size (Å) | 1.223 | 1.223 |
| Symmetry imposed | C1 | C1 |
| Initial particle images (no.) | 362,740 | 51,588 |
| Final particle images (no.) | 38,278 | 17,707 |
| Map resolution (Å) | 7.3 Å | 8.3 Å |
| FSC threshold | 0.143 | 0.5 |
| **Refinement** |  |  |
| Initial model used | 5FD7, 6TZ9, AF-Q8NC56-F1, | AF-Q8NC56-F1 |
| (PDB or AlphaFold code) | AF-Q8WUX9-F1, |  |
| Q-score | 0.22 | N/A |
| Map sharpening B factor (Å<sup>2</sup>) | -10 | N/A |
| **Validation** |  |  |
| MolProbity score | 0.63 | N/A |
| Clashscore | 0.35 |  |
| Poor rotamers (%) | 0.18 |  |
| **Ramachandran plot** |  |  |
| Favored (%) | 98.02 | N/A |
| Allowed (%) | 1.82 |  |
| Disallowed (%) | 0.16 |  |

### Molecular dynamics simulations

#### Structural models

First, we constructed MD models from a 111-subunit ring model wherein every subunit constitutes a CHMP1B-CHMP7-LEM2 protomer (Supp. Fig. 9). Intrinsically disordered C-terminal regions were removed to define constructs: 1) CHMP1B residues 1–130, 2) CHMP7 residues 1–365, and 3) full-length LEM2 residues 1–503. Therefore, filament assemblies retained core structural integrity while excluding disordered termini.

In addition, the microtubule lattice was built from the microtubule cryo-EM structure (PDB 3JAT (61,62)), comprising 14 protofilaments and 5 dimer layers. We adjusted the lattice edges to ensure compatibility with the simulation box along z, producing a dimer rise of 8.4 nm and a microtubule continuous across the periodic boundary.

Furthermore, conformational ensembles of the LEM2 low-complexity domain (LCD, residues 71–201) were generated via hierarchical chain growth (63), assembling full-length disordered chains from exhaustively sampled all-atom fragment libraries. We grew 500 chains, utilizing 42 for LCD-only slab simulations and 96 for the MT-LCD system. Two coarse-grained carbon nanotubes (mean radius 7.4 nm, length 3.7 nm) were parameterised (64) and included in the flat region of the nuclear-envelope membrane systems. Each nanotube punctures one bilayer of the double membrane only to allow water exchange with the lumen.

#### All-atom simulations

We employed the CHARMM36m protein force field (65,66) and CHARMM36 lipids (67) for membrane systems, alongside the TIP3P water model (68,69) all systems were solvated and neutralised at 0.15 M NaCl. Individual CHMP7 (open and closed conformations) and CHMP1B (open conformation) monomers were placed in cubic boxes for two independent 1 µs replicates. In addition, the membrane-embedded CHMP1B–CHMP7–LEM2 trimer was constructed by back-mapping an equilibrated coarse-grained configuration (97.5 ns) using CG2AT2 (70). The membrane reproduced an endoplasmic-reticulum/nuclear-envelope-like composition (30 % DOPC, 30 % POPC, 10 % POPE, 10 % DOPE, 10 % POPI, 3 % POPS, 2 % DOPS, 1 % POPA, 1 % DOPA, and 3 % cholesterol) at an area per lipid of 0.6 nm². The system was placed in a triclinic box (47.2 x 47.2 x 29.8 nm), and water molecules within 4 Å of any lipid were deleted. This system was simulated in three independent 1 µs replicates, differing only in the random seed for initial velocities. Neighbour lists used the Verlet scheme (1.4 nm cut-off, updated every 20 steps), while electrostatics were treated with particle-mesh Ewald (PME); real-space Coulomb and Lennard-Jones cut-offs were 1.2 nm, and periodic boundary conditions were applied in all three dimensions. Each system was energy-minimised by steepest descent to a maximum force below 10 kJ mol⁻¹ nm⁻¹, followed by NVT equilibration (375 ps, 1 fs time step) and NPT equilibration (2 ns, 2 fs time step). Temperature was held at 310 K with the velocity-rescaling thermostat (0.01 ps coupling during NVT, 0.1 ps subsequently), applied separately to protein and non-protein particles; pressure was held at 1 bar with the Parrinello–Rahman barostat (2.0 ps coupling, 4.5 × 10⁻⁵ bar⁻¹ compressibility), using isotropic coupling for monomers and semiisotropic coupling for the membrane trimer. Altogether, production runs were unrestrained, used a 2 fs time step, with coordinates written every 1 ns.

#### Coarse-grained simulations

We employed the Martini 2.2 force field (71,72) with non-polarisable water (10 % antifreeze, ε = 15). Proteins were mapped to coarse-grained resolution using martinize.py (73) with the ElNeDyn elastic-network model (74) and DSSP 3.1.4 (75,76) secondary structure assignment. To mitigate systematic over-attraction (77), we rescaled selected protein–protein Lennard-Jones well depths using the martiscale utility (32). Specifically, the LEM2-LCD region (residues 1–200) was assigned to a bead class with a 0.70 scaling factor (tuned from slab phase separation simulations), while interactions between this region and the remainder of the system were scaled by 0.84; all other protein–protein interactions remained unscaled.

In addition, we constructed nuclear-envelope pore systems using a flat coarse-grained bilayer (10-component composition) generated with insane.py (78). A double membrane traversed by a cylindrical hole was generated with BUMPy 1.1.0 (79), and two carbon nanotubes were inserted into an additional 8.5 nm hole in the flat region, following the same protocol reported for NPC simulations (80). Four distinct systems were built: 1) protein-free pore; 2) LEM2-only filament; 3) CHMP7-LEM2 filament; and 4) full CHMP1B–CHMP7–LEM2 filament (incomplete ring, 82 subunits, box 40 x 140 x 42 nm, from Supp. Fig. 9). To emulate microtubules crossing the pore, we extended the lattice to 7 dimer layers and inserted three periodic copies at the pore center. Furthermore, we characterised LEM2 LCD-MT interactions by combining a tail-less microtubule with 96 LCD chains in a triclinic box (80 x 80 x 42 nm).

Systems were solvated and 0.15 M NaCl was added. Water and ions within 4 Å of protein or lipid were removed. Equilibration proceeded in three stages: 1) 1 ns NVT (5 fs time step); 2) 2.5 ns NPT (5 fs time step, Berendsen barostat, backbone restraints); and 3) 97.5 ns NPT (15 fs time step, Berendsen barostat, backbone restraints). Production runs were unrestrained, used a 15 fs time step, and employed the Parrinello–Rahman barostat (81). Temperature was maintained at 310 K (velocity rescaling (82), 1.0 ps coupling), and pressure at 1 bar (semiisotropic coupling, 12.0 ps coupling). Electrostatics used a reaction field (**ε** = 15) with 1.2 nm cut-offs. Each system was simulated in three independent replicates (2 µs for pore/MT systems, 20 µs for MT-LCD only system).

All simulations, and all preparation steps requiring MD simulation software, used GROMACS 2024 (83). System assembly, clash detection, leaflet assignment and analysis were implemented in Python with MDAnalysis (84) and Biopython (85).

### Microtubule binding assays

#### In vitro reconstitution and imaging

All microtubule binding assays were performed at room temperature. Microtubule filaments (labeled with Rhodamine-tubulin) were grown from TAMRA labeled tubulin monomers in BRB80 buffer as described above. For the reconstitution, two coverslips (22 x 22 mm^2^ and 18 x 18 mm^2^; Corning, Inc., Corning, NY), were cleaned in piranha solution (H_2_O_2_/H_2_SO_4_, 3:5; both Sigma) before silanization with 0.05% dichlorodimethylsilane in trichloroethylene (Sigma) and connected by heating pieces of Parafilm M (Pechiney Plastic Packaging, Chicago, IL). to form 3 flow channels (86). These channels were coated with anti-beta-tubulin antibodies (0.2 mg/ml in BRB80, Sigma). The solution containing microtubule filaments were then passed through the channels to allow for immobilization of microtubules. Unbound filaments were then washed out with BRB80 buffer. They were inspected for stable localizations using an inverted TIRF microscope (Axio Observer Z1; Carl Zeiss Microscopy GmbH) with a 63× oil immersion objective (NA 1.46; Zeiss) with an additional 1.6x optovar lens. For binding assays with LEM2_Lem-LCD_, varying concentrations of LEM2_Lem-LCD_ (500nM, 1 µM or 5 µM) were added. LEM2_Lem-LCD_ was labeled with DyLight-650 (1:100 stoichiometry of labeled to unlabeled). For binding assays with spastin, spastin-GFP (Δ227 isoform) (produced in the Diez lab (87) was added at a concentration of 20 nM. The laser lines (Omicron Laserbox Sole-3) and filtersets used were 488 nm (BP482/18, zt491, BP520/35), 561 nm (ex 520/35, em 585/40, dc 532) and 647 nm (ex 628/40, em 692/40, dc 635) respectively. For timelapse imaging, snapshots were taken at an interval of 20 sec for 15 min using a EMCCD detector (iXon Ultra; Andor Technology) and MetaMorph software controle (Molecular Devices Corporation).

#### Image analysis

All raw image files were annotated with acquisition metadata (imaging channel, timepoint, experimental condition, and replicate identifier) to generate a curated, traceable dataset. Representative fields of view were assembled into multi-panel montages for figure preparation and visual quality control prior to quantitative analysis.

#### Motion correction

Time-lapse image stacks were corrected for sample drift using high-intensity fiducial markers present in the field of view (tubulin speckles). For each frame, the displacement relative to a reference frame (frame 1) was estimated from the fiducial positions and expressed as a per-frame position transformation matrix. This transformation was applied uniformly to all imaging channels to preserve inter-channel spatial registration. Motion-corrected image stacks were saved as TIFF files and stored in a dedicated directory for downstream processing.

### Visualization

Figures were prepared using Chimera 1.16, ChimeraX 1.8. and 1.10.1, 3dmod from the IMOD package, Adobe Illustrator, or Microsoft Excel. The membrane model shown in Fig. 6 was generated in Blender 3.5.0 (http://www.blender.org).

## Notes

### Competing Interest Statement

The authors have declared no competing interest.

